# Genetic architecture of soluble arabinoxylan fibre in elite genotypes of bread wheat revealed by genome-wide association analysis

**DOI:** 10.64898/2026.07.21.739768

**Authors:** Abdul Kader Alabdullah, Ondrej Kosik, Michelle Leverington-Waite, Rowan A.C. Mitchell, Anneke Prins, James Brett, Simon Griffiths, Peter R. Shewry, Alison Lovegrove

## Abstract

Dietary fibre intake remains below recommended levels, and increasing fibre content of widely consumed white wheat flour (derived from the starchy endosperm) represents a scalable strategy to improve public health. In wheat starchy endosperm, arabinoxylan (AX) is the dominant fibre component, with the water-extractable (WE) fraction being particularly beneficial to health. We assembled an Elite Fibre Panel (EFP) of 384 elite modern wheat genotypes from UK commercial breeding programmes and quantified the content of WE-AX in wholemeal, as a proxy for soluble AX in white flour, across two UK field environments. Wholemeal WE-AX, showed substantial quantitative variation and moderate genotype-by-environment interaction, with a broad-sense heritability of 0.68. Genome-wide association analyses using 6,791 SNPs identified seven loci associated with WE-AX content. The strongest and most stable effects were mapped to major loci on chromosomes 1B and 6B, previously implicated in AX regulation, while additional loci of smaller and sometimes environment-dependent effect were detected on chromosomes 3A, 5B, and 7A. Favourable alleles increased WE-AX content by ∼5-15% and combined additively. LD-defined intervals contained several high-confidence candidate genes involved in cell-wall biosynthesis, remodelling and post-depositional modification, including PER1, a validated regulator of arabinoxylan cross-linking, together with genes encoding a UTP-glucose-1-phosphate uridylyltransferase, trichome birefringence-like proteins and xyloglucan endotransglucosylase/hydrolases. These findings demonstrate that substantial gains in soluble AX can be achieved by pyramiding favourable alleles already segregating within elite germplasm, providing a practical route for breeding wheat with enhanced dietary fibre content and improved nutritional quality.

**Key message:** Multiple additive loci controlling soluble arabinoxylan content were identified in elite wheat germplasm, enabling marker-assisted breeding for increased dietary fibre in white flour.

## Introduction

Dietary fibre (DF) is a key component of a healthy human diet, with fibre intake being associated with reduced risk of major non-communicable diseases, including type 2 diabetes, cardiovascular disease, and colorectal cancer. Furthermore, large prospective cohort studies and meta-analyses have, over a number of years, demonstrated inverse associations between total dietary fibre intake and all-cause mortality, as well as disease-specific outcomes (Aune et al., 2016; Veronese et al., 2018; Reynolds et al., 2019; Oh et al., 2019; Barber et al., 2020; Dahm et al., 2024). These health benefits have led to DF being recognised as a critical target for public health policies worldwide.

Dietary fibre comprises a heterogeneous group of carbohydrate oligomers and polymers and associated compounds that resist digestion in the upper gastrointestinal (GI) tract (Jones, 2014). Major DF components include non-starch (cell wall) polysaccharides, fructo-oligosaccharides, and resistant starch, with the precise components, their proportions and their physicochemical properties varying widely among food sources. The beneficial effects of DF result from multiple, complementary mechanisms operating along the GI tract. In the upper GI tract, fibre can modulate the rate of food breakdown, digestion, and nutrient absorption, thereby attenuating postprandial glycaemic responses. In the colon, fermentable fibres are metabolised by the gut microbiota to produce short-chain fatty acids (SCFAs), such as acetate, propionate, and butyrate, which support colonocyte health and exert systemic metabolic effects after absorption. In addition, fibre increases faecal bulk and reduces intestinal transit time, limiting exposure of the gut epithelium to potentially harmful compounds (Gill et al., 2021; Reynolds et al., 2019).

The physiological behaviour of DF in the GI tract is strongly influenced by three interrelated properties: solubility, viscosity, and fermentability. Soluble fibre tends to increase the viscosity of digesta, slowing starch digestion and glucose absorption in the small intestine and reducing glycaemic load, while also being more rapidly fermented in the proximal colon. By contrast, insoluble fibre is fermented more slowly, often in the distal colon, and contributes more to faecal bulk. It is increasingly recognised that a balanced intake of different fibre types is required to maximise health benefits across the GI tract (Gill et al., 2021; Reynolds et al., 2026).

Despite the well-established health benefits of DF, fibre intake remains substantially below recommended levels in many populations. Governmental and regulatory bodies typically recommend adult fibre intakes of 25-30 g per day. However, average intakes are considerably lower in many countries: approximately 16-17 g/day in UK adults compared with a target of 30 g/day, and around 16 g/day in the United States against a similar recommendation, with little or no change in intake over the last 40 years (NDNS, 2025; US Department of Health and Human Services, 2020; Derbyshire, 2026). These persistent shortfalls highlight the need for population-level strategies to increase fibre intake without requiring major changes in dietary habits.

Cereal-based foods, particularly wheat products, represent one of the most important contributors to dietary fibre intake in industrialised diets. In the UK, bread alone provides approximately 12% of total fibre intake in adults (Bates et al., 2025). Wholegrain wheat products are rich in fibre, with wholemeal bread typically containing 7g fibre/100g compared with about 2.9g/100g in white bread (McCance and Widdowson’s composition of foods integrated dataset, 2021). However, although the consumption of wholegrain products has been promoted for several decades, the majority of wheat-based foods consumed in the UK and globally are produced from white flour rather than wholemeal flour, and the proportion of wholemeal and brown flours milled in the UK has declined in recent years (Shewry et al., 2023).

White flour is derived almost exclusively from the starchy endosperm, the major storage tissue of the wheat grain, which accounts for approximately 82% of grain dry weight (Barron et al., 2007). During milling, the fibre-rich bran layers and embryo are removed, resulting in a substantially lower fibre content. Typically, white flour contains around 4% DF (dry weight), of which approximately half is arabinoxylan (AX), a cell wall polysaccharide which is sometimes referred to as pentosan (arabinose and xylose being pentose sugars) (Shewry et al., 2024). Increasing fibre content in white flour, and in particular AX, is therefore an attractive strategy to enhance fibre intake at the population level, as it targets widely consumed foods without increasing cost or requiring changes in consumer behaviour.

The AX content of white flour varies approximately two-fold, from about 1.35% to 2.7% (dry weight), with genetic factors accounting for 50-75% of the observed variation and the remainder attributable to environmental effects and genotype × environment interactions (Gebruers et al., 2008; Martinant et al., 1988; Shewry et al., 2010; Tremmel-Bede et al., 2020; Hernandez-Espinosa et al., 2020). This substantial heritable variation provides a strong foundation for genetic improvement.

Genetic analyses using biparental mapping populations initially identified three meta-QTL influencing total and/or soluble AX content in white flour, located on chromosomes 1B, 3D, and 6B. These loci were also detected through association analysis in a small diversity panel of approximately 150 lines (Quraishi et al., 2011). The chromosome 1B locus has since been consistently validated as a major determinant of both total (TOT) and soluble (water-extractable, WE) AX content through analyses of biparental populations derived from crosses involving the high-AX cultivar Yumai 34, as well as through broader association studies (Lovegrove et al., 2020; Ibba et al., 2021). By contrast, the locus on chromosome 6B primarily affects soluble AX content and has been shown to correspond to a defective cell wall peroxidase gene involved in oxidative cross-linking of AX via ferulate dimerisation (Lovegrove et al., 2020; Mitchell et al., 2026). More recently, Jiang et al. (2024) identified additional loci for AX content through association analysis of 205 wheat lines, including loci specific to soluble and insoluble AX fractions.

Although several studies have investigated the genetic control of WE-AX in wholemeal flour, total AX measurements from wholemeal are not directly informative for *white flour* because the total content of AX is much higher in bran than in white flour, by about x10, most of which is insoluble (Gebruers et al., 2008). However, the contents of WE-AX in wholemeal and white flours are strongly correlated (r = 0.96) (Shewry et al., 2024). The content of WE-AX in wholemeal can therefore be used as a proxy for WE-AX in white flour in large-scale screening and genetic analyses, increasing the throughput by avoiding the requirement for milling to produce white flour (Lovegrove et al., 2020). Consistent with this, QTL for WE-AX in wholemeal flour have been reported on chromosomes including 1B and 6B, though not consistently on 3D (Yang et al., 2016; Li et al., 2024; Chen et al., 2024).

In this study, we assembled an Elite Fibre Panel (EFP) comprising 384 elite modern wheat genotypes, including advanced breeding lines and released cultivars provided by commercial breeding programmes in the UK. Field trials conducted on two sites revealed substantial variation in WE-AX in wholemeal flour and genome-wide association analysis (GWAS) was used to determine the genetic basis of this variation and the extent to which previously characterised AX-associated loci are represented in modern wheat. Our analyses confirmed the presence of the major loci on chromosomes 1B and 6B within the panel and revealed additional loci of smaller effects, indicating that WE-AX content is controlled by a combination of major and polygenic components in elite backgrounds. Importantly, favourable alleles were present but not fixed across loci, highlighting the potential for further improvement through targeted selection.

The presence of high-AX alleles in elite, high-performing breeding lines and cultivars demonstrates that increased fibre content is compatible with good agronomic performance and end-use quality. Together, these findings establish the Elite Fibre Panel as a powerful resource for dissecting the genetic architecture of dietary fibre in wheat and support the feasibility of improving fibre content in white flour by exploiting allelic variation within modern breeding programmes, without the requirement to introgress variation from exotic germplasm.

## Materials and Methods

### Plant Materials

#### A) Elite Fibre Panel (EFP)

The Elite Fibre Panel (EFP) comprised 384 elite wheat breeding lines and cultivars supplied by four commercial breeding companies: RAGT Ltd., Saaten Union, Limagrain UK Ltd., and F1 Seeds Ltd. Of these, 365 lines were retained for analysis, with 19 excluded due to incomplete phenotypic or genotypic data. The panel represents germplasm from the United Kingdom, France, and Germany (**Table 1, Table S1**) and provides a diverse, commercially relevant resource for GWAS of wheat grain fibre traits.

**Table 1.**
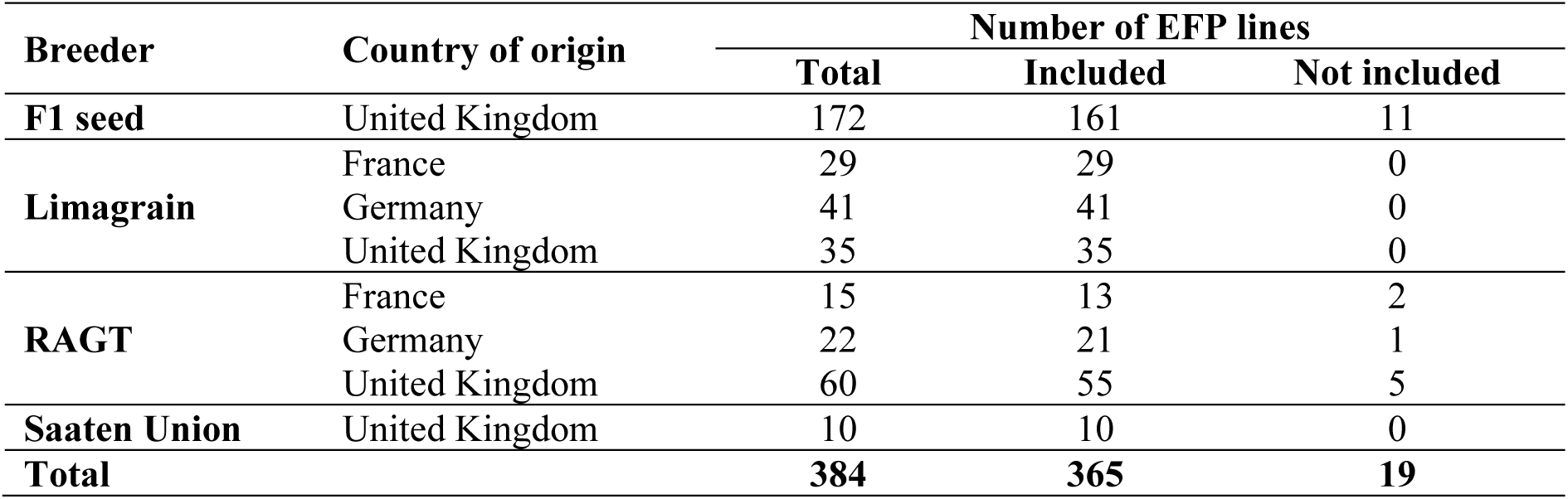
The number of Elite Fibre Panel (EFP) lines from each breeder and country of origin.

### Field trials

All EFP genotypes were grown in 1 x 2 metre plots on two sites, RAGT Ltd. (Ickleton, Essex, UK) and Limagrain Ltd. (Woolpit, Suffolk, UK) in the season 2022-2023. Plots were harvested using small plot combine harvesters, where the yield per plot and the grain moisture were measured.

### Wholemeal arabinoxylan analysis

Wholemeal flour was produced from mature grain using a Retsch centrifugal mill (250 µm sieve) and stored at −20 °C until analysis. WE-AX in aqueous extracts of wholemeal flour was determined using the semi-high throughput method described by Hernández-Espinosa et al., 2024 (a modified phloroglucinol colorimetric pentosan assay for the quantification of arabinoxylan) and expressed as xylose equivalents (**Table S2**).

### DNA extraction and genotyping

Young leaf tissue from each EFP line was collected at the 2-3 leaf stage and stored at −80 °C prior to DNA extraction. Genomic DNA was isolated from ∼50 mg tissue using the LGC Oktopure system. Genome-wide SNP genotyping was performed with the Axiom TaNG1.1 array (44,266 SNPs), and genotype calling was conducted using the Axiom™ Analysis Suite (AxAS v5.3.0.45) under default settings. Targeted genotyping of known fibre-related QTLs was carried out using KASP assays (**Table S3**) following standard protocols, with genotype clusters analysed using KlusterCaller.

### Genome-wide Association Analysis (GWAS)

GWAS was conducted following stringent SNP quality control, in which monomorphic markers and those with >10% missing data or >10% heterozygosity were excluded (**Table S4**). The remaining markers were analysed using the GAPIT3 (v.3.1.0) R package (Wang and Zhang, 2021), with a minor allele frequency threshold of 0.05 applied to retain informative SNPs. Population structure and pairwise genetic relatedness were estimated within GAPIT using principal component analysis (PCA) and the Zhang kinship algorithm, respectively. These parameters were incorporated into the GWAS models according to their respective statistical frameworks. PCA was performed using eigenvalue decomposition, and the appropriate number of principal components to include as covariates was selected empirically by comparing genomic inflation factors (λ) and the number of significant marker-trait associations across models fitted with 0-10 principal components.

To assess GWAS model performance, a simulation-based power analysis implemented in GAPIT was carried out using simulated phenotypes derived from the observed genotype data (heritability h² = 0.9; number of quantitative trait nucleotides = 5). Multiple GWAS models (GLM, MLM, MLMM, CMLM, ECMLM, FarmCPU, BLINK, and SUPER) were evaluated across 100 simulation replicates, and their power, false discovery rate, and type I error were compared. Based on this evaluation, the three best-performing models were selected for final analyses. Genome-wide association analyses were then performed using the optimal number of PCs, with Manhattan and quantile-quantile plots generated using the rMVP package. Based on the effective number of independent markers (M_eff_ = 6,791), a Bonferroni-corrected genome-wide significance threshold of -log₁₀(p) ≥ 5.02 (p ≤ 9.57 × 10⁻⁶) was applied. Significant marker trait associations (MTAs) located within 1 Mb on the same chromosome were grouped and considered to represent a single genomic locus. For each locus, the MTA showing the strongest statistical support was designated as the lead SNP and used to define the LD-based QTL interval for subsequent candidate gene identification.

### LD-based SNP-specific determination of QTL intervals

Genome-wide LD patterns were initially examined using GAPIT to characterise marker correlations, inter-marker distances and the overall relationship between LD and physical distance across the EFP genome. Chromosome-wide pairwise LD matrices and LD-decay profiles were subsequently calculated in R using the *snpStats* package (Clayton, 2015), with mean *R*^2^summarised in adaptive physical-distance bins containing a target minimum of 30 SNP pairs, to assess chromosome-specific variation in LD structure and decay. For each SNP significantly associated with a trait (MTA), a local linkage disequilibrium (LD) analysis was performed to define the corresponding QTL interval. All SNPs located on the same chromosome as the lead SNP were extracted, ordered by physical position based on the wheat reference genome assembly IWGSC v1, and LD (R²) between the lead SNP and neighbouring markers were calculated using the *snpStats* package. The primary LD threshold used to delineate the QTL interval was R² ≥ 0.5, and the threshold was also relaxed to R² ≥ 0.3 to ensure that a biologically meaningful interval was defined. In each case, the genomic region spanned by all SNPs exceeding this threshold was taken as the QTL boundary. In cases where no such supporting SNPs were identified, the interval was collapsed to the physical position of the lead SNP. This LD-based, marker-specific approach allowed data-driven determination of QTL intervals that reflect local haplotype structure and recombination patterns.

### Candidate Gene Identification

Candidate genes were identified by extracting all annotated genes located within the linkage disequilibrium (LD)-defined interval surrounding each significant SNP from the wheat reference genome (IWGSC RefSeq v1.1) using EnsemblPlants. Candidate genes were prioritised using a multi-criteria approach that integrated genomic position, predicted gene function, developmental expression profiles, and published biological evidence. Functional annotations were examined to identify genes with plausible roles in cell wall biosynthesis and remodelling, arabinoxylan metabolism, carbohydrate metabolism, nucleotide-sugar biosynthesis and transport, oxidative cross-linking, and grain development. Gene expression during grain development was assessed using publicly available starchy endosperm RNA-seq datasets available through the Wheat Expression Browser (Borrill *et al*., 2016; Ramírez-González *et al*., 2018) and the Kino *et al*. (2020) study (**Table S8**), with particular emphasis on genes expressed during the period of active endosperm cell-wall formation. Additional supporting evidence was incorporated where available by assessing the evolutionary conservation of candidate genes using estimates of monocot-, commelinid-, and grass-specificity derived from the Universal Grass Gene Database (Mitchell, 2025). The highest-priority candidate gene for each QTL was selected based on the overall weight of evidence rather than genomic position alone.

### High-Fibre Allele Frequency

High-Fibre Allele (HFA) frequency in the EFP was calculated for each MTA and compared with that obtained from the variant matrix of 827 Watkins landrace and 224 wheat varieties (Cheng et al., 2024). Accessions with missing or heterozygous SNP calls were not considered for allele frequency calculation.

## Results

### Phenotypic variation and heritability of soluble arabinoxylan content

The content of WE-AX in wholemeal flour showed extensive phenotypic variation across the EFP when evaluated at two independent UK field trial sites. Across environments, WE-AX content spanned almost a three-fold range, from approximately 3.1 to 8.7 mg g⁻¹ at the Limagrain site and 2.7 to 8.2 mg g⁻¹ at the RAGT site (**Fig. 1a; Table S2**), demonstrating substantial exploitable variation within elite modern wheat germplasm.

**Fig. 1.**
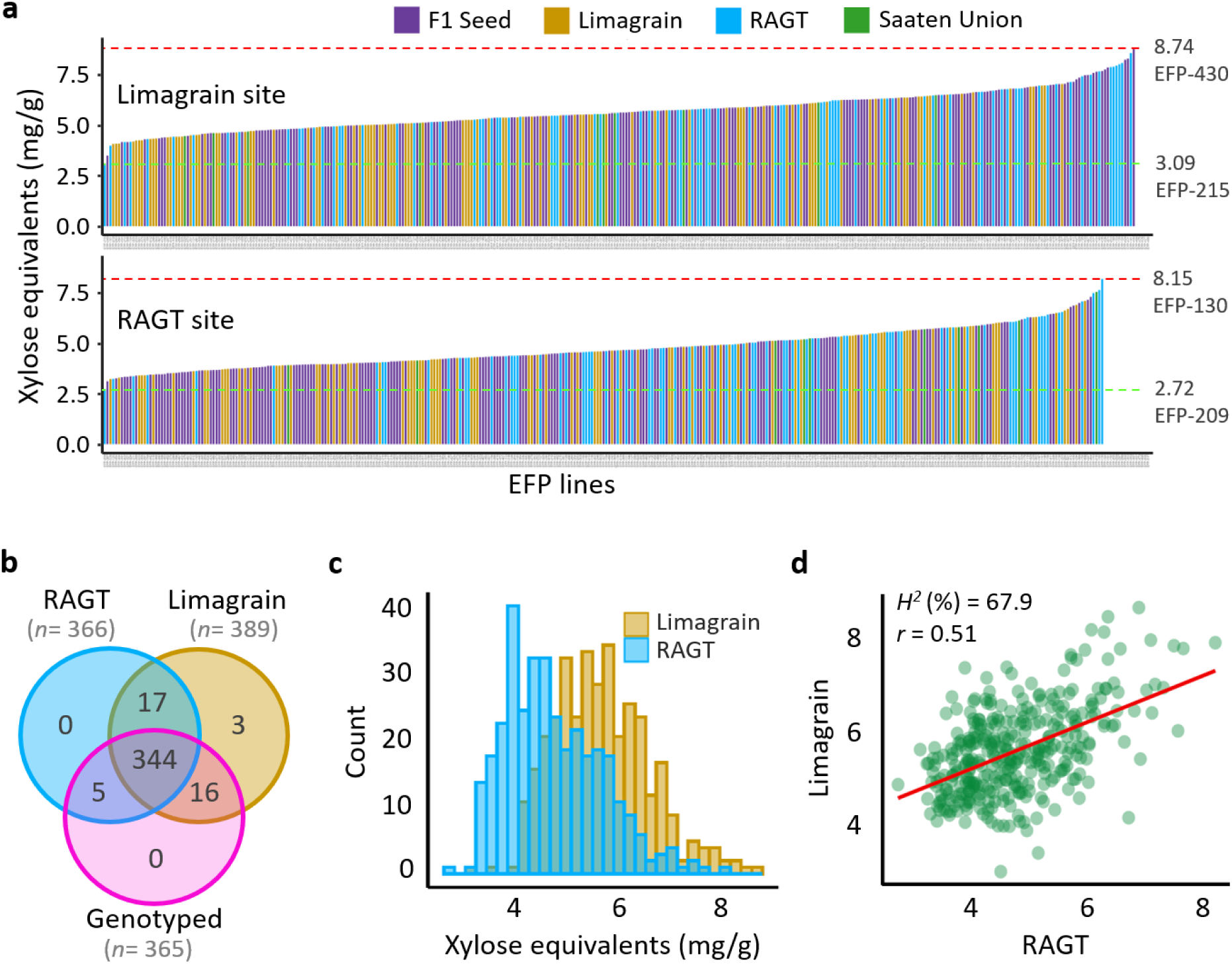
Summary of the Elite Fibre Panel (EFP) used in this study. **a)** Ranked distribution of WE-AX content in wholemeal (expressed as xylose equivalents) for individual EFP genotypes grown at the Limagrain (upper panel) and RAGT (lower panel) field trial sites. Genotypes are ordered from low to high xylose content within each site, and bars are coloured according to breeder origin. Dashed horizontal lines indicate the minimum and maximum observed values at each site, with selected extreme genotypes highlighted. **b)** Venn diagram illustrating the overlap of EFP lines that were genotyped and phenotyped across the three datasets. The majority of lines (344) were included in all datasets. **c)** Distribution of the WE-AX content (expressed as xylose equivalents, in mg g⁻¹) in wholemeal across EFP lines grown in field trials conducted by Limagrain and RAGT. Histograms show overlapping frequency distributions for each site, highlighting differences in trait variation between the two environments. Number of bins = 30. **d)** Relationship between WE-AX (expressed as xylose equivalents) of genotypes grown at the Limagrain and RAGT sites. Each point represents the WE-AX contents in wholemeal of individual genotypes, with the solid line indicating the linear regression fit. Broad-sense heritability (H^2^) and the Pearson correlation between genotypes across the two sites (r) are shown.

The overlap of genotyped and phenotyped lines across datasets is shown in **Fig. 1b**, with the majority of genotypes (344 lines) common to all datasets, providing a robust basis for downstream analyses. The distribution of WE-AX values was continuous at both sites, consistent with quantitative inheritance. Frequency distributions of xylose content differed between the two environments (**Fig. 1c**), with the Limagrain site displaying a higher mean WE-AX content (5.67 mg g⁻¹) than the RAGT site (4.73 mg g⁻¹), indicating a significant environmental effect on trait expression. Phenotypic variability also differed between sites, with coefficients of variation (CV) of 16.4% at Limagrain and 20.0% at RAGT, indicating greater dispersion in genotype performance at the latter environment.

However, variance partitioning revealed a relatively high broad-sense heritability (H² = 0.68), highlighting a strong genetic contribution to water soluble AX accumulation across environments (**Fig. 1d**). Genotypic performance showed a moderate but significant correlation between sites (r = 0.51; *P* < 0.001), suggesting that while shared genetic determinants influence WE-AX content, genotype-by-environment interactions also play an important role (**Fig. 1d**). This is consistent with the estimate of heritability. Together, these results indicate that selection for increased soluble AX is feasible within elite breeding material, although environmental context should be considered during phenotypic evaluation.

### Genome-wide association analysis identifies seven loci associated with WE-AX content

A total of 6,791 high-quality SNP markers were retained following stringent quality control, which excluded monomorphic markers and those with >10% missing data or >10% heterozygosity (**Tables S4** and **S5**; **Fig. S1**). Before genome-wide association analysis, the genetic structure of the Elite Fibre Panel was evaluated using principal component analysis (PCA). Principal component analysis revealed only moderate population structure within the EFP. The first three principal components explained 13.78% of the total genetic variation, and no discrete genetic clusters were observed (**Fig. 2a-c**). To identify the most appropriate GWAS approach, the performance of seven statistical models implemented in GAPIT was compared using simulation analyses. Based on 100 simulation replicates, the analyses showed that the models MLMM, FarmCPU, and BLINK consistently provided the highest statistical power while maintaining effective control of false-positive associations (**Fig. 2d, e**). The influence of population structure correction was then assessed by fitting models with 0 - 10 principal components. Across all three phenotypic datasets (Average, Limagrain, and RAGT), genomic inflation factors (λ) remained close to the expected value of one irrespective of the number of principal components included, indicating that population structure and relatedness were adequately controlled without additional PCA correction (**Fig. 2f; Fig. S2a-c**). Likewise, the number of genome-wide significant marker-trait associations detected by the three best-performing models remained largely unchanged across the range of principal component corrections evaluated (**Fig. 2g; Fig. S2d-f**). Together, these results demonstrate that inclusion of principal components had little influence on either model calibration or the detection of significant associations. Consequently, BLINK, FarmCPU, and MLMM were selected for the final GWAS analyses, which were performed without principal component correction (PCA.total = 0). Consistent with the PCA results, the genomic kinship matrix showed generally low pairwise relatedness across the EFP, although several small groups of more closely related accessions were evident (**Fig. S3**).

**Fig. 2.**
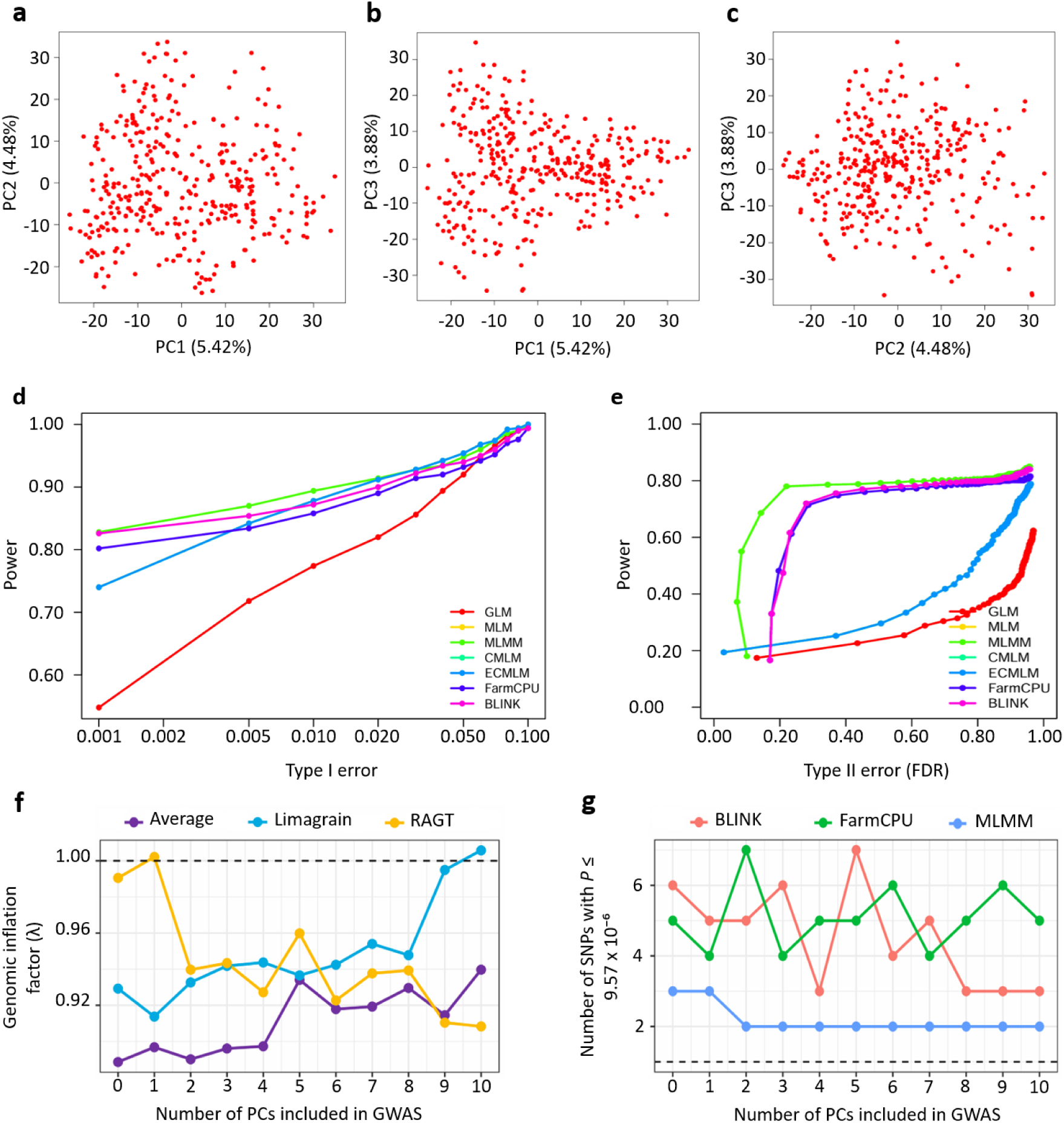
Population structure of the Elite Fibre Panel (EFP), evaluation of principal component correction, and comparison of GWAS model performance. Principal component analysis (PCA) was performed using 6,791 high-quality SNP markers from 365 wheat accessions. Scatterplots showing the genetic relationships among accessions based on (**a**) PC1 versus PC2, (**b**) PC1 versus PC3, and (**c**) PC2 versus PC3. The percentage of genetic variation explained by each principal component (Eigenvalue Percentage; EVP) is indicated on the corresponding axes. Panels (d) and (e) compare the statistical performance of seven GWAS models implemented in GAPIT (GLM, MLM, MLMM, CMLM, ECMLM, FarmCPU, and BLINK) based on 100 simulation replicates. (**d**) Statistical power plotted against the Type I error rate and (**e**) statistical power plotted against the false discovery rate (FDR). (**f**) Genomic inflation factor (λ) estimated using the MLM model following inclusion of 0-10 principal components for the Average, Limagrain, and RAGT phenotypic datasets. The dashed line indicates the expected value under the null hypothesis (λ = 1). (**g**) Number of genome-wide significant associations (P ≤ 9.57 × 10⁻⁶) detected using BLINK, FarmCPU, and MLMM for the average WE-AX phenotype following inclusion of 0-10 principal components.

GWAS were conducted using 6,791 high-quality SNPs and the three best-performing statistical models (BLINK, FarmCPU, and MLMM) to ensure the robust detection of marker-trait associations. A Bonferroni-corrected genome-wide significance threshold of -log₁₀(p) ≥ 5.02 (p ≤ 9.57 × 10⁻⁶) was used based on M_eff_ = 6,791. Seven loci significantly associated with WE-AX content were identified across environments and analytical models (**Fig. 3a**). Quantile-quantile (Q-Q) plots indicate a high quality of the detected genome-wide associations using the three statistical models (**Fig. 3b**).

**Fig. 3.**
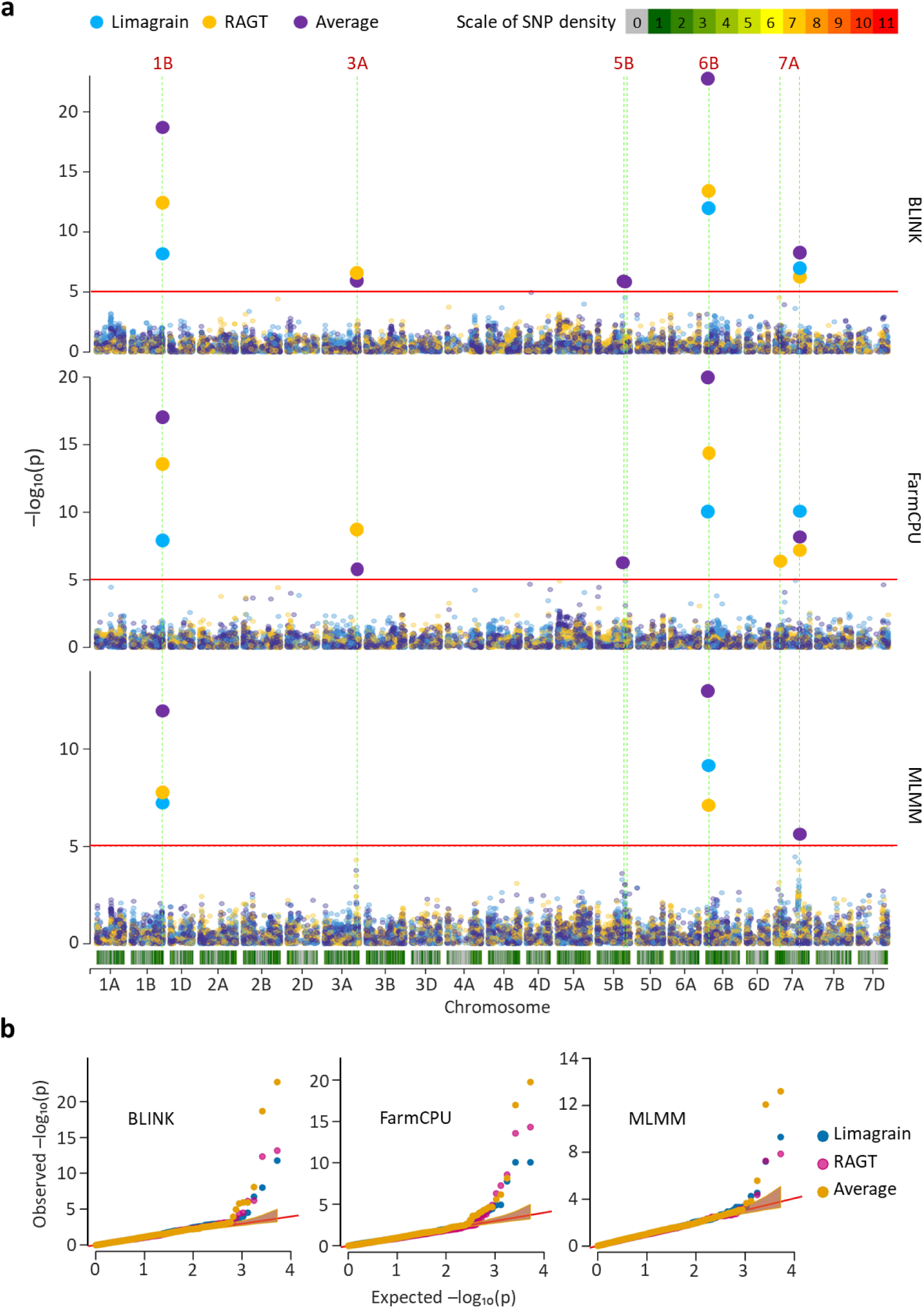
GWAS of WE-AX content (expressed as xylose equivalents) in wholemeal flours of the Elite Fibre Panel. **(a)** Manhattan plots showing GWAS results using three multi-locus models: BLINK (top), FarmCPU (middle), and MLMM (bottom). Analyses were performed using phenotypic data from the Limagrain site, the RAGT site, and the mean across both sites. The horizontal red line denotes the genome-wide significance threshold (-log₁₀ *p* = 5.02). Vertical dashed green lines indicate chromosomes harbouring significant association peaks. The bars beneath each Manhattan plot represent SNP density calculated in 1-Mb sliding windows, with marker counts shown according to the colour scale (ranging between 0-11 SNPs per Mb). **(b)** Quantile-quantile (Q-Q) plots for each GWAS model and trait, comparing observed versus expected -log₁₀(*p*) values. These plots illustrate deviations from the null hypothesis and demonstrate close agreement between observed and expected values for most markers, with deviations in the upper tail corresponding to significant associations.

The most significant associations were detected on chromosomes 1B and 6B, corresponding to previously characterised major QTLs for arabinoxylan content (Lovegrove et al., 2020; Ibba et al., 2021; Mitchell et al., 2026). The lead SNP on chromosome 1B (JB1B_662960818; a KASP marker) exhibited strong and consistent effects across all environments and models (-log₁₀(p) = 13.58), confirming the importance of this locus in controlling soluble AX levels in elite germplasm. Similarly, the locus on chromosome 6B (a KASP marker for PER1) showed the strongest statistical support observed in this study (-log₁₀(p) = 22.74), consistent with the finding that this SNP directly influences AX solubility through modulation of cell-wall cross-linking (Mitchell et al., 2026). Another significant association on chromosome 7A was detected constantly across environments and statistical models, where the lead SNP was AX-643812441 (-log₁₀(p) = 10.08; **Table 2**).

**Table 2.** Significant SNPs associated with WE-AX (expressed as xylose equivalents) in wholemeal identified by GWAS across environments and models.

| MTA | Chr | Position | $-\log_{10}(p)$ | FDR | Effect | Model | Trait |
| --- | --- | --- | --- | --- | --- | --- | --- |
| JB1B_662960818 | 1B | 652451624 | 13.58 | 6.83E-11 | -0.54 | All | All |
| AX-643813054 | 3A | 682369519 | 8.58 | 4.61E-06 | 0.26 | BLINK,<br>FarmCPU | RAGT,<br>Average |
| AX-94909517 | 5B | 536591354 | 6.19 | 8.40E-04 | -0.24 | BLINK,<br>FarmCPU | Average |
| AX-95679056 | 5B | 600496907 | 5.80 | 1.36E-03 | 0.38 | BLINK | Average |
| PER1 | 6B | 26076727 | 22.74 | 9.57E-20 | 0.23 | All | All |
| AX-643821508 | 7A | 88621297 | 6.33 | 4.85E-04 | 0.27 | FarmCPU | RAGT |
| AX-643812441 | 7A | 511923724 | 10.08 | 2.19E-07 | -0.25 | All | All |

*FDR* represents the false discovery rate-adjusted p-value, used to account for multiple testing across the genome. *FDR* was calculated according to the Benjamini-Hochberg procedure (Benjamini & Hochberg, 1995). *Effect* denotes the estimated allelic effect on xylose content, with the sign indicating the direction of the effect (positive values indicate an increase and negative values a decrease in xylose content associated with the alternative allele). The terms $-\log_{10}(p)$ , *FDR*, and *Effect* are reported for the trait-model combination in which the MTA exhibited the strongest statistical support (i.e. the largest $-\log_{10}(p)$ ). The designation “All” indicates that the corresponding MTA was detected as significant across all traits or across all GWAS models evaluated in this study.

In addition to these three loci, four additional associations were detected on chromosomes 3A, 5B, and 7A (**Fig. 3a**). These loci generally exhibited smaller effect sizes and, in some cases, environment-specific significance, suggesting more context-dependent contributions to WE-AX variation. Collectively, the seven loci explain a substantial proportion of the genetic variance for soluble AX content observed in the EFP. A detailed list of all detected MTAs is provided in **Table S6**.

### Allelic effects of significant loci for grain WE-AX content identified by GWAS

Allelic substitution analysis at the seven significant loci revealed consistent and biologically meaningful effects on wholemeal WE-AX content (**Fig. 4**). For each locus, clear differences in WE-AX content were observed between homozygous allelic classes, with the favourable (high-fibre) allele conferring increases of approximately 5-15% relative to the alternative allele. These effects were generally stable across environments, being evident in the across-environment mean as well as in the individual Limagrain and RAGT field trials. In most cases, allelic differences were highly significant (ANOVA, *p* < 0.001), and *post hoc* comparisons confirmed statistically distinct allele classes based on Tukey’s HSD test. A small number of loci displayed reduced or non-significant effects in one environment, indicating a degree of environmental modulation, but the overall direction of effect remained consistent. Collectively, these results demonstrate that the identified loci exert additive and robust contributions to variation in WE-AX content in elite wheat germplasm and highlight their potential value for marker-assisted improvement of grain fibre content.

**Fig. 4.**
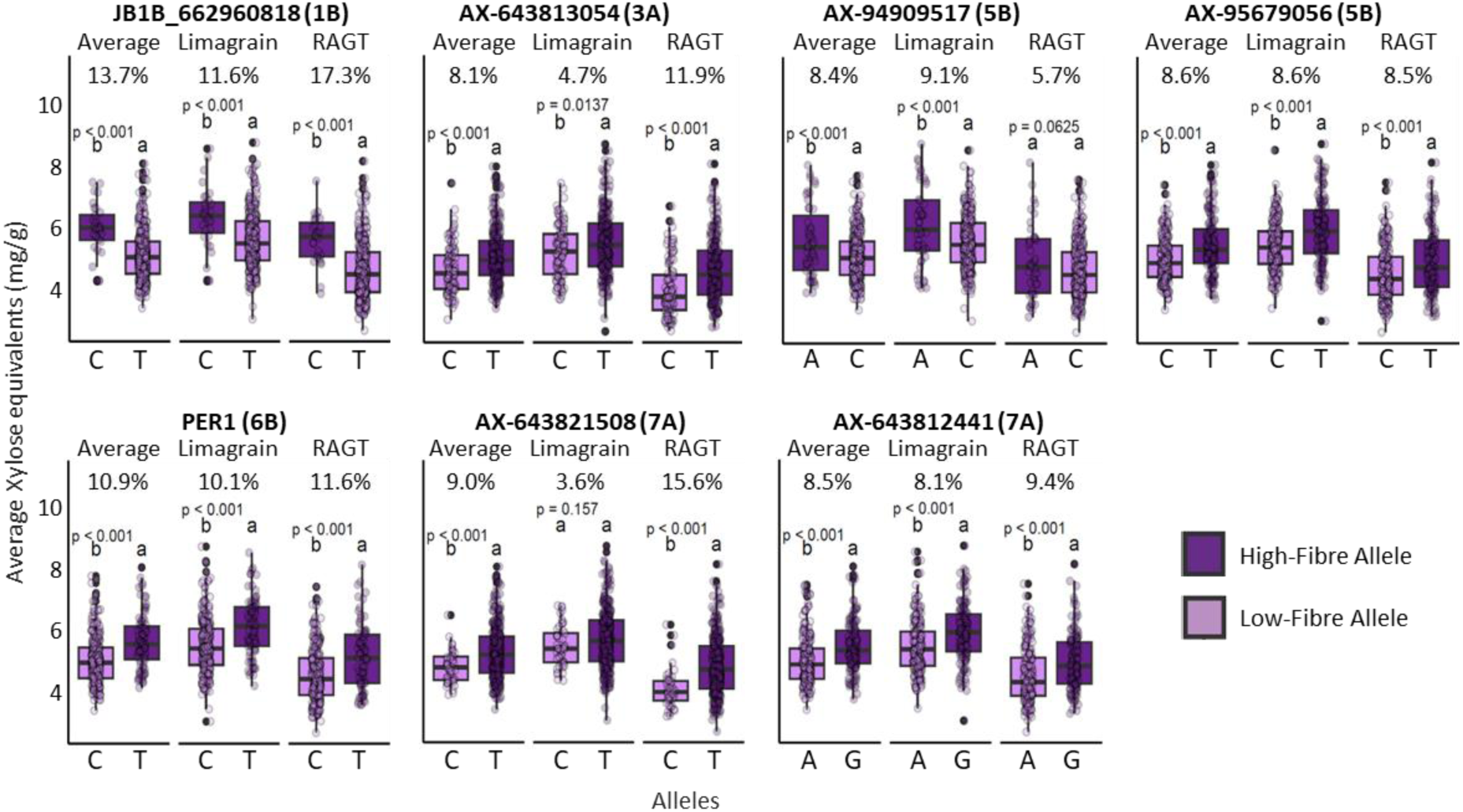
Allelic effects of MTAs on wholemeal flour WE-AX content (expressed as xylose equivalents) across environments. Boxplots show the distribution of WE-AX content (mg g⁻¹) for homozygous alleles at the seven MTAs, analysed across three datasets: the across-environment mean (*Average*), and individual field trials conducted by Limagrain and RAGT. Only homozygous genotypes were included in the analysis. The percentage values above each panel indicate the relative difference in mean xylose content between the two alleles, calculated as the percentage increase of the higher-mean allele relative to the lower-mean allele. Statistical significance of allelic differences within each dataset was assessed using one-way analysis of variance (ANOVA). Different letters above boxplots denote statistically significant differences between alleles based on Tukey’s honestly significant difference (HSD) post hoc test (p < 0.05). ANOVA p-values are shown above each dataset panel. Chromosome assignments for each MTA are indicated in parentheses next to the marker ID. Dots represent individual observations.

### Frequency of high-fibre alleles in the elite fibre panel

**Fig. 5a** summarises the distribution of high- and low-fibre alleles at the seven genome-wide significant loci within the EFP. The frequency of the high-fibre allele varied markedly among markers, indicating substantial heterogeneity in the representation of favourable alleles across elite germplasm. At several loci, including 6B (PER1), the high-fibre allele occurred at intermediate to high frequencies, suggesting that these favourable alleles are already incorporated into modern breeding material. In contrast, loci on chromosomes 1B (JB1B_662960818) and 5B (AX-94909517) showed relatively low frequencies of the high-fibre alleles, indicating considerable scope for further improvement through targeted selection. Comparison with a broader collection of modern varieties and Watkins landraces showed substantial differences in favourable-allele frequencies among genetic resources (**Fig. S4**; **Table S7**), reflecting their contrasting breeding histories and genetic diversity. Together, these findings demonstrate that, although several major fibre-enhancing alleles are already widely represented in elite wheat, others remain underutilised and provide valuable opportunities to increase flour WE-AX through allele pyramiding in breeding programmes.

**Fig. 5.**
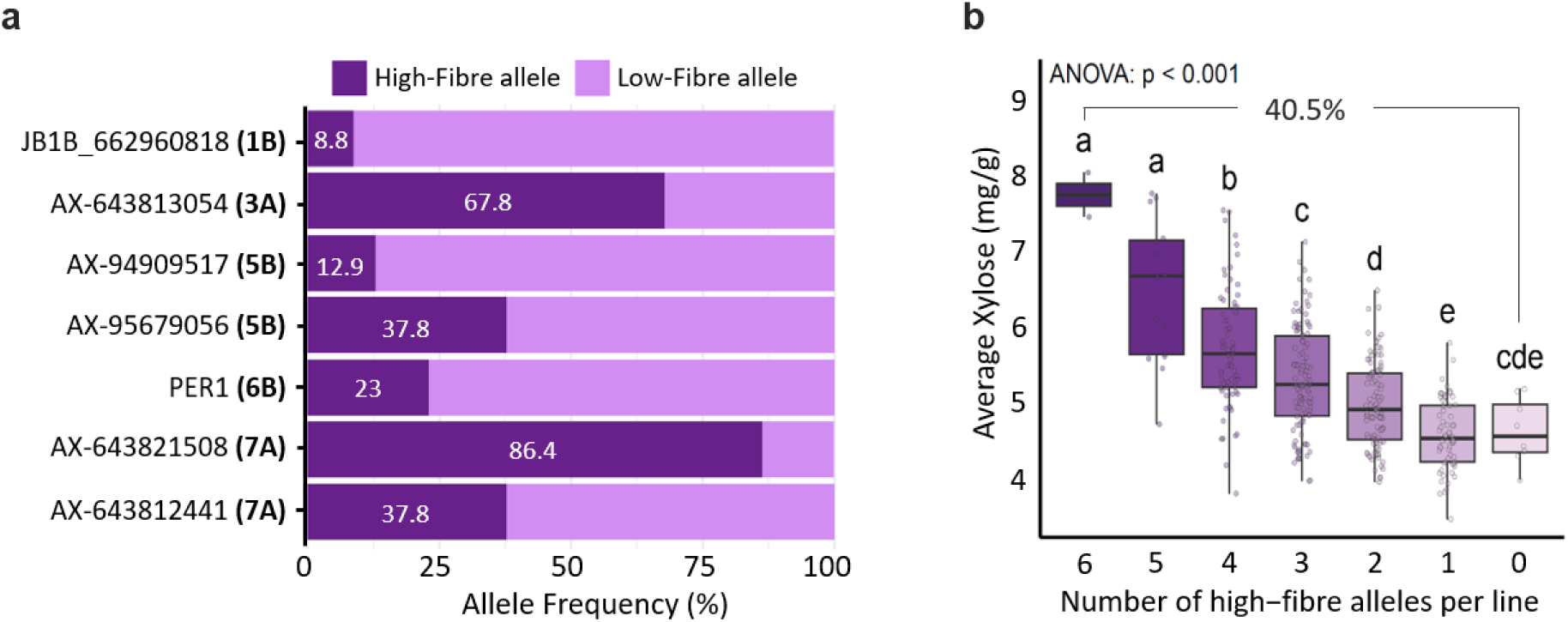
High-fibre allele frequencies and their cumulative effect on wholemeal WE-AX content (expressed as xylose equivalents) in the EFP. (**a**) Stacked bar plots showing the frequency (%) of the high-fibre allele (dark purple) and the low-fibre allele (light purple) at seven significant GWAS-associated SNP markers (JB1B_662960818, AX-643813054, AX-94909517, AX-95679056, PER1, AX-643821508, and AX-643812441) across the Elite Fibre Panel. Allele frequencies were calculated from genotyping matrix of the EFP. Values shown on bars indicate the frequency (%) of the high-fibre allele. (**b**) Boxplots illustrating the relationship between average grain WE-AX content (mg g⁻¹) and the cumulative number of high-fibre alleles carried across the seven GWAS-identified SNPs. Each box represents the distribution of mean WE-AX values for lines carrying between 0 and 6 high-fibre alleles. Different letters denote statistically significant differences among allele-count classes based on one-way ANOVA followed by Tukey’s HSD post hoc test (*p* < 0.001). The percentage value indicates the relative difference in mean WE-AX content between lines carrying no high-fibre alleles and those carrying the maximum number of favourable alleles.

### Cumulative contribution of high-fibre alleles to WE-AX variation in the Elite Fibre Panel

Analysis of the cumulative number of favourable alleles across the seven significant loci revealed strong and largely additive effects on grain WE-AX content (**Fig. 5b**). Lines carrying progressively higher numbers of high-fibre alleles exhibited stepwise increases in average WE-AX content, with statistically significant differences observed between most allele-count classes (one-way ANOVA, *p* < 0.001). In particular, genotypes carrying six favourable alleles - the maximum observed within the Elite Fibre Panel - showed substantially higher WE-AX levels compared with lines lacking favourable alleles, corresponding to an overall increase of approximately 40% in mean WE-AX content. No genotype carried all seven favourable alleles, indicating that complete pyramiding of the identified loci has not yet been achieved in elite germplasm. Together, these findings demonstrate that the effects of individual loci are largely additive and suggest that further increases in grain WE-AX content can be achieved by stacking existing favourable alleles within elite germplasm, reducing the need to introduce variation from exotic germplasm.

### Linkage disequilibrium-defined QTL intervals and candidate gene content

Genome-wide LD analysis indicated an overall decline in marker association with increasing physical distance, while chromosome-specific analyses revealed substantial variation in LD structure among chromosomes and genomic regions (**Figs. S5-S7**). This heterogeneity supported the use of marker-specific local LD patterns, rather than a single genome-wide distance threshold, to delineate the QTL intervals shown in **Fig. 6**. Linkage disequilibrium (LD)-based interval mapping revealed substantial variation in the physical extent of the detected QTL regions, ranging from very narrow intervals (<1 Mb) containing one or a few annotated genes to broad regions spanning several tens of megabases and encompassing hundreds of genes (**Fig. 6**). Notably, the locus on chromosome 6B (PER1) was defined by a particularly narrow LD interval, consistent with strong local LD and supporting the genotyped SNP as the actual cause of this association and acting as a perfect marker since it leads to a mislocalisation of the encoded peroxidase (Mitchell et al., 2026). Investigating the gene content of the LD-defined intervals of the WE-AX QTLs reveals a total of 725 high-confidence genes, out of which 414 genes were expressed in starchy endosperm (**Fig. 6; Table S9**).

**Fig. 6.**
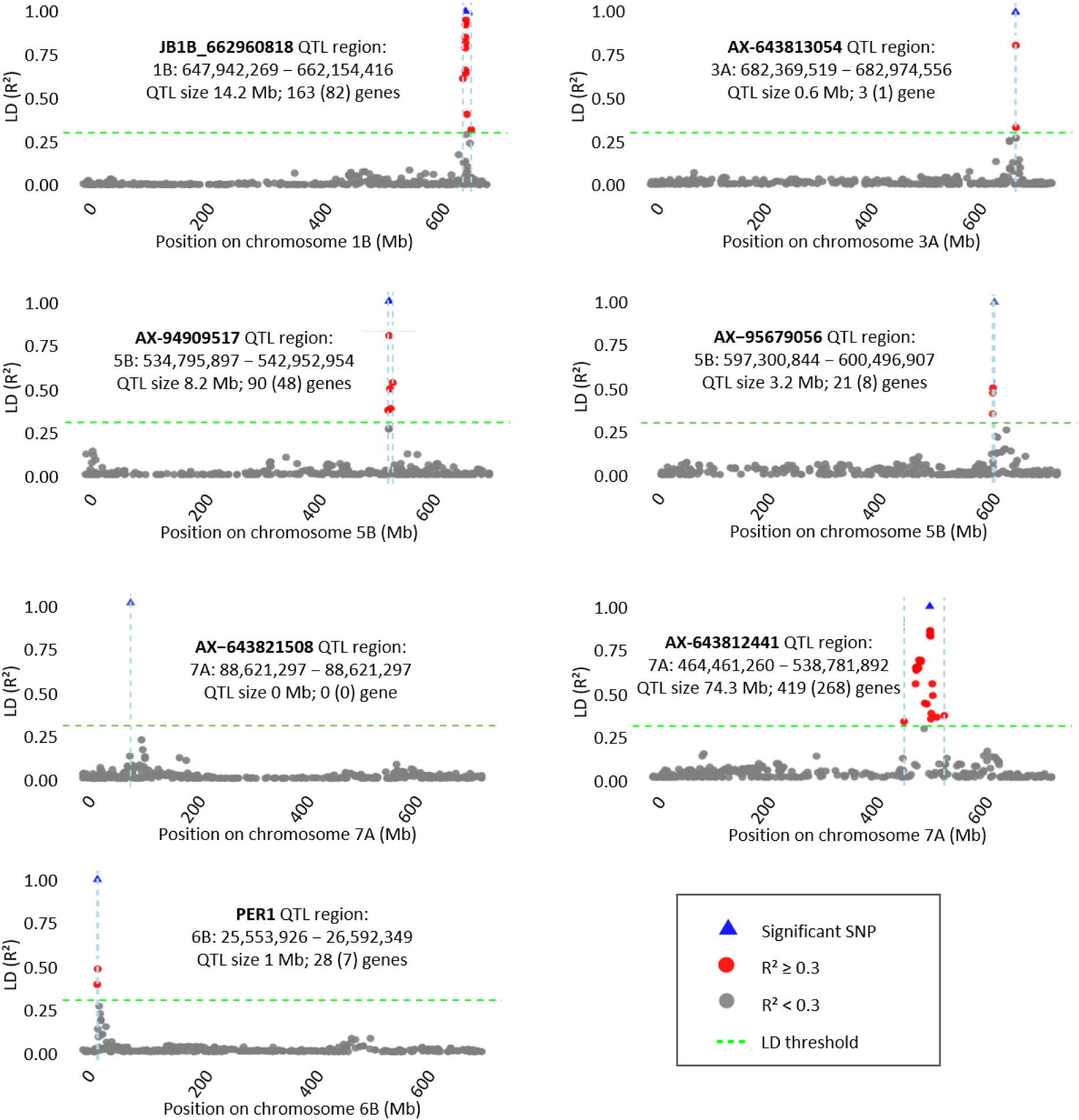
LD-based delineation of QTL intervals for SNPs significantly associated with WE-AX content of wholemeal flour. Local linkage disequilibrium (LD) patterns surrounding the lead SNPs were used to define quantitative trait locus (QTL) intervals for wholemeal flour WE-AX content. For each genome-wide significant SNP, LD (R²) was calculated between the lead SNP (blue triangle) and all neighbouring SNPs on the same chromosome, ordered by physical position. Points represent individual SNPs coloured by LD strength relative to the lead SNP (red, R² ≥ 0.3; grey, R² < 0.3). The vertical dashed blue line indicates the physical position of the lead SNP, while horizontal dashed green lines denote the LD threshold used to define QTL boundaries. The genomic coordinates shown within each panel correspond to the final LD-defined QTL regions on chromosomes 1B, 3A, 5B, 6B, and 7A. The number of genes shown in parentheses indicates the number of genes expressed in the endosperm (TPM ≥ 1 in at least one sample). For the marker AX-643821508 on chromosome 7A, no neighbouring markers exceeded the LD threshold; the QTL interval collapsed to the physical position of the lead SNP.

### Candidate gene identification

Candidate-gene analysis identified several biologically plausible genes within the LD-defined intervals of the WE-AX QTLs (**Table 3; Table S9**). Candidates were prioritised by integrating functional annotation, expression in developing starchy endosperm, evolutionary specificity and evidence from experimentally characterised genes or orthologues. The chromosome 6B interval provided the strongest functional evidence, containing PER1 (*TraesCS6B02G042500*) and the adjacent peroxidase gene *TraesCS6B02G042600*. PER1 is a commelinid-specific class III cell-wall peroxidase for which a defective allele has been experimentally shown to reduce ferulate dimerisation and AX cross-linking, thereby increasing the water-extractable AX fraction; this establishes *TraesCS6B02G042500* as the causal or near-causal gene underlying the 6B association (Mitchell et al., 2026). Both genes were strongly expressed in developing endosperm and classified as commelinid-specific in the universal_grass_peps resource, consistent with a specialised role in grass cell walls, in which feruloylated arabinoxylan and ferulate-mediated cross-links are characteristic features (Mitchell, 2025).

**Table 3.**
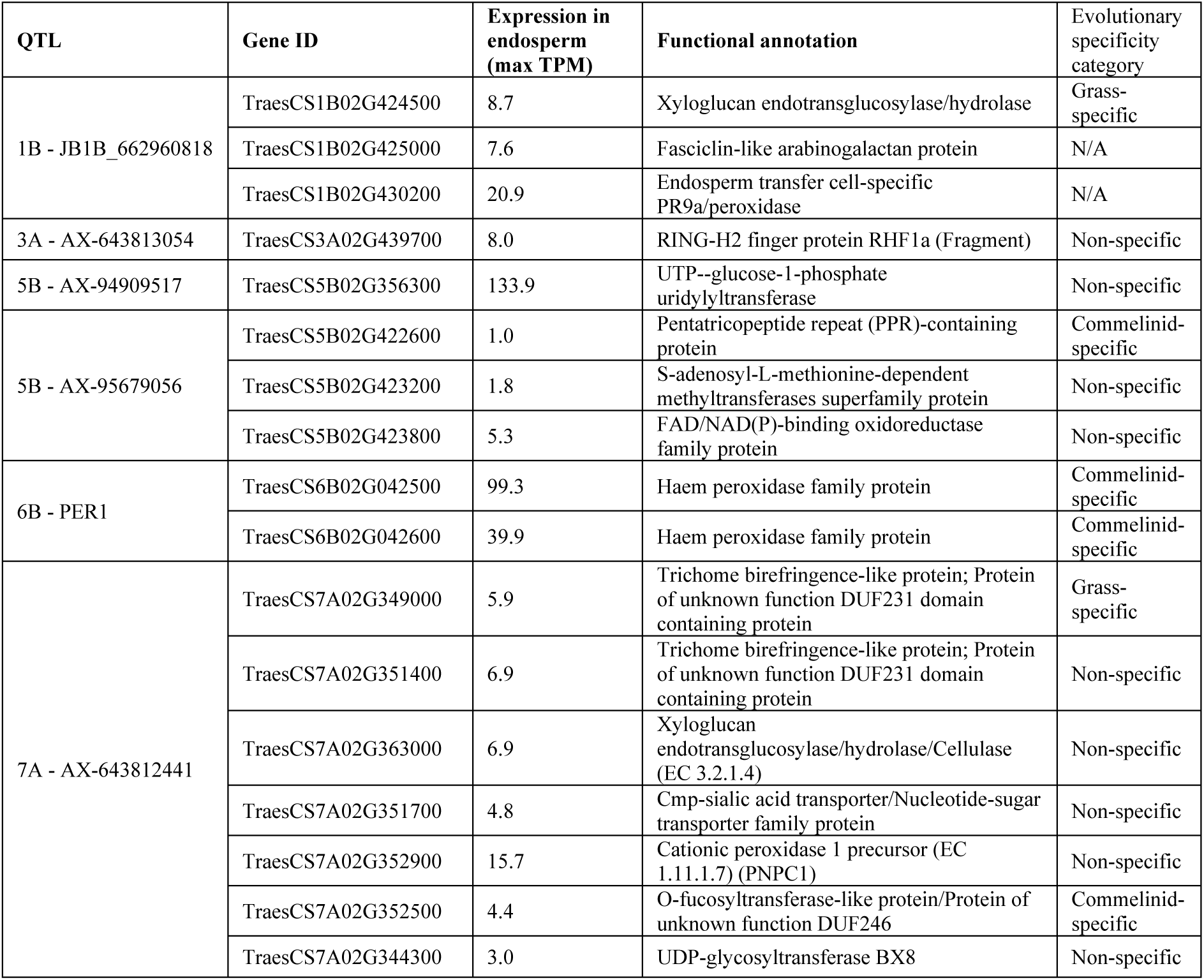
Significant GWAS-associated SNPs for grain WE-AX content and candidate genes within LD-defined QTL intervals.

The major chromosome 1B interval contained three particularly plausible candidates. The first, *TraesCS1B02G424500*, encodes a grass-specific xyloglucan endotransglucosylase/hydrolase (XTH/GH16). XTH proteins cleave and reconnect xyloglucan chains and are established regulators of cell-wall assembly, loosening and remodelling, although studies in rice indicate that many XTHs act principally on xyloglucan rather than directly on arabinoxylan; its candidacy should therefore be interpreted as an effect on overall wall architecture and polysaccharide organisation rather than direct AX synthesis (Van Sandt, et al., 2007). A second candidate, *TraesCS1B02G425000*, encodes a fasciclin-like arabinogalactan protein, a class of extracellular glycoproteins associated with cell-wall organisation, cellulose deposition and mechanical properties; notably, fasciclin-like arabinogalactan proteins have been detected in the wheat-grain cell-wall proteome (Liu, et al., 2020). The third, *TraesCS1B02G430200*, encodes an endosperm transfer-cell-specific PR9a peroxidase and is therefore mechanistically plausible because class III peroxidases can modify cell-wall phenolics and promote oxidative cross-linking. However, unlike PER1, no direct biochemical or genetic evidence currently links this particular peroxidase to AX solubility.

At the chromosome 5B QTL marked by AX-94909517, the strongest candidate was *TraesCS5B02G356300*, encoding UTP-glucose-1-phosphate uridylyltransferase (UGPase). UGPase catalyses the formation of UDP-glucose, a central precursor in plant carbohydrate metabolism that feeds the production of UDP-glucuronic acid and, subsequently, UDP-xylose and UDP-arabinose, the activated sugar donors required for xylan and arabinoxylan biosynthesis (Kleczkowski et al., 2004). Its very high expression in developing endosperm further supports a role in supplying nucleotide-sugar substrates during grain cell-wall deposition. By contrast, the candidates at the second 5B locus are less directly connected to arabinoxylan biosynthesis but remain biologically plausible. The commelinid-specific pentatricopeptide repeat (PPR)-containing protein *TraesCS5B02G422600* represents the strongest candidate at this locus. Although PPR proteins are primarily recognised as regulators of organellar RNA processing, translation and RNA editing in mitochondria and plastids (Barkan and Small, 2014), increasing evidence indicates that specific PPR proteins also contribute to cell wall integrity and maintenance. For example, the Arabidopsis Cell Wall Maintenance 1 (CWM1) and CWM2 PPR proteins function as sensors of cell wall integrity, linking defects in organellar gene expression to adaptive responses that maintain cell wall homeostasis under stress (Hofmann, 2016; Hu et al., 2016). Consequently, *TraesCS5B02G422600* may influence WE-AX indirectly through regulatory mechanisms affecting cell wall development or maintenance rather than by participating directly in arabinoxylan biosynthesis. The remaining candidates at this locus include an S-adenosyl-L-methionine (SAM)-dependent methyltransferase and an FAD/NAD(P)-binding oxidoreductase. SAM-dependent methyltransferases constitute a large enzyme family involved in the methylation of diverse metabolites, including cell-wall-associated phenylpropanoids, lignin precursors and other secondary metabolites, whereas FAD/NAD(P)-binding oxidoreductases participate in numerous oxidation-reduction reactions associated with primary metabolism, reactive oxygen species homeostasis and cell-wall metabolism (Joshi and Chiang, 1998; Vanholme, et al., 2019; Mhamdi et al., 2010). Although these functions make both genes biologically plausible candidates, there is currently no direct experimental evidence linking either gene family to arabinoxylan biosynthesis or water-extractable arabinoxylan accumulation in wheat.

The chromosome 7A interval contained several candidates related to hemicellulose synthesis or modification. Of these, *TraesCS7A02G349000* and *TraesCS7A02G351400* encode trichome birefringence-like proteins containing DUF231 domains. Members of the TBL/DUF231 family have been biochemically demonstrated to act as polysaccharide O-acetyltransferases, including xylan O-acetyltransferases in both Arabidopsis and rice, making these genes strong candidates for modifying xylan structure, wall interactions and extractability (Gao, et al., 2017). The grass-specific classification of *TraesCS7A02G349000* provides additional evolutionary support for a specialised function in grass cell-wall metabolism (Mitchell, 2025). Another strong pathway-level candidate, *TraesCS7A02G351700*, encodes a nucleotide-sugar transporter-family protein. Golgi-localised nucleotide-sugar transporters supply activated sugars for matrix-polysaccharide biosynthesis, and experimentally characterised UDP-xylose transporters are required for normal xylan content and structure (Zhao, et al., 2018). Additional candidates included an XTH, a cationic peroxidase, an O-fucosyltransferase-like protein and a UDP-glycosyltransferase, although their links to AX remain less direct.

The chromosome 3A LD interval contained a single annotated gene, TraesCS3A02G439700, encoding a RING/U-box protein (Table 3). RING/U-box proteins function primarily as E3 ubiquitin ligases that regulate protein turnover and numerous developmental and stress-response pathways in plants. TraesCS3A02G439700 exhibited its highest expression at 4 dpa, followed by a gradual decline throughout grain development (Fig. 7), suggesting a role during early endosperm differentiation. Although no direct evidence currently links this gene family to arabinoxylan biosynthesis or modification, its unique presence within the LD interval makes it the primary candidate gene underlying this QTL.

**Fig. 7.**
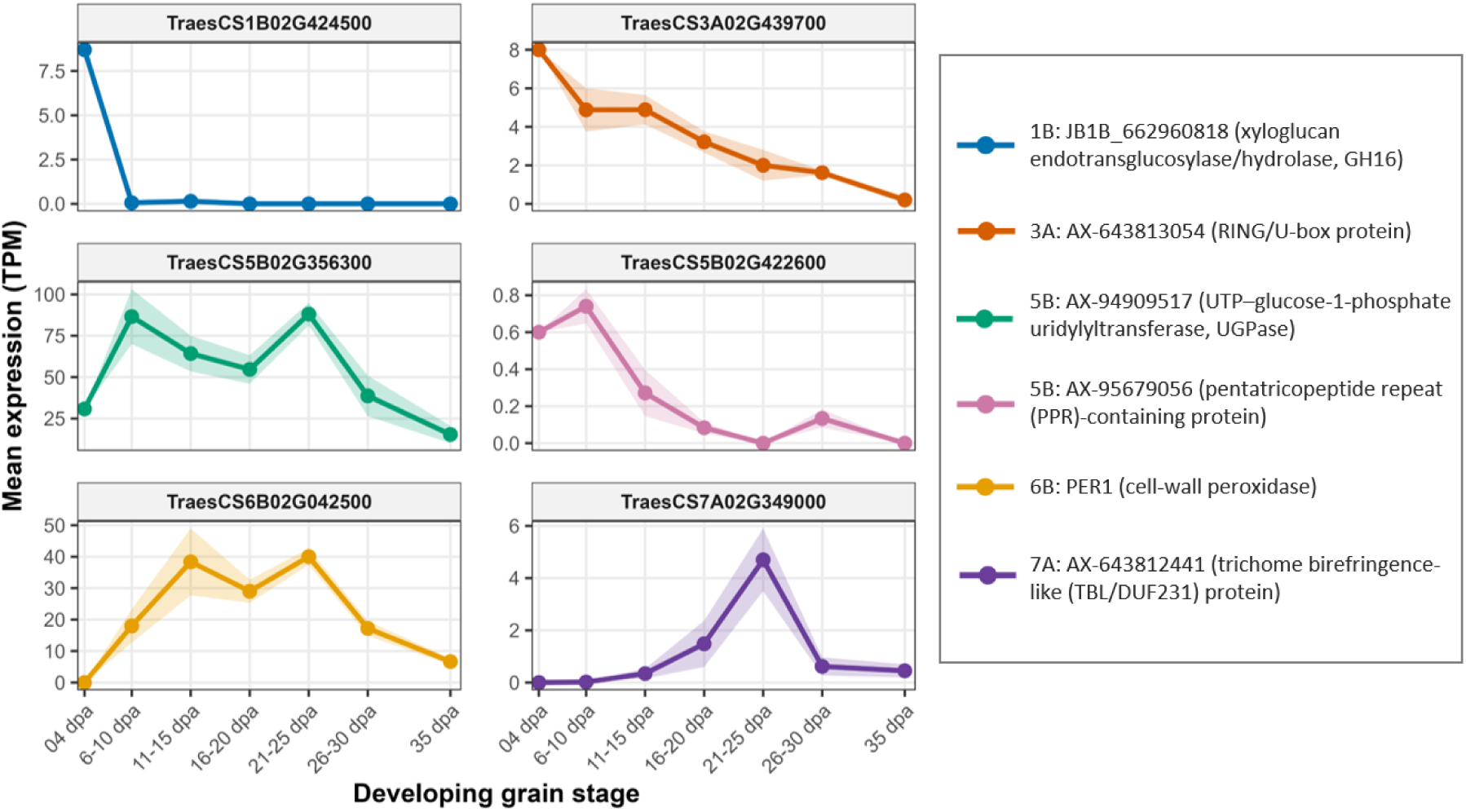
Endosperm developmental expression profiles of the highest-priority candidate genes underlying the major water-extractable arabinoxylan (WE-AX) QTLs identified by genome-wide association analysis. Gene expression data were obtained from developing starchy endosperm sampled at seven developmental stages (4-35 days post anthesis; dpa). The dataset comprises 29 RNA-seq samples previously reported by Ramírez-González *et al*. (2018) (17 samples) and Kino *et al*. (2020) (12 samples) (see Table S8 for details). Gene expression abundance is expressed as transcripts per million (TPM). Lines represent the mean expression across biological replicates, and shaded ribbons indicate ± standard error (SE). The diverse temporal expression profiles suggest that these candidate genes may function at different stages of endosperm development, with several exhibiting peak expression during the period of active cell-wall biosynthesis and maturation.

The second chromosome 7A locus (AX-643821508) did not contain any candidate genes because no neighbouring SNPs were in significant linkage disequilibrium with the lead SNP (**Fig. 6**). Consequently, the LD-defined interval collapsed to 0 Mb, preventing the assignment of a genomic interval for candidate gene identification.

Many of the prioritised genes were expressed in developing starchy endosperm and displayed distinct temporal profiles, with UGPase, PER1, the 1B PR9a peroxidase and the 7A TBL gene reaching their highest expression at stages consistent with active cell-wall construction or maturation (Fig. 7; Table 3). Collectively, the evidence identifies PER1 as the best-supported causal gene, while the 1B XTH, 5B UGPase and 7A TBL and nucleotide-sugar transporter genes represent the strongest novel functional candidates. Nevertheless, except for PER1, these assignments remain hypotheses that require confirmation through allele-specific expression, variant-effect analysis, fine mapping and functional validation.

## Discussion

This study provides new insights into the genetic architecture of water-extractable arabinoxylan (WE-AX) content in white flour from elite European bread wheat genotypes by combining genome-wide association analysis with linkage disequilibrium (LD)-based interval mapping, developmental transcriptomics, evolutionary conservation, and functional annotation. Seven loci associated with WE-AX content were identified, including the previously reported major loci on chromosomes 1B and 6B together with four additional loci on chromosomes 3A, 5B and 7A. Importantly, the favourable alleles identified at these loci remain segregating within elite breeding germplasm rather than being fixed, demonstrating that substantial genetic variation for increasing dietary fibre already exists within modern breeding material. The persistence of these favourable alleles despite intensive selection for high yield, disease resistance and adaptation indicates that enhanced fibre content is compatible with modern breeding objectives and can be further improved through selection within elite breeding populations.

The chromosome 6B locus containing PER1 represents the strongest and best-supported association identified in this study. This locus produced the most significant association across all GWAS models and environments and was defined by a very narrow LD interval, consistent with the causal variant residing within the lead marker. Recent functional studies demonstrated that PER1 encodes a defective class III cell-wall peroxidase that reduces ferulate dimerisation and oxidative cross-linking of arabinoxylan, thereby increasing the water-extractable fraction without substantially altering total arabinoxylan content (Mitchell et al., 2026). This mechanism provides a direct biological explanation for the large effects on WE-AX observed in the present study, while also illustrating that natural variation in soluble fibre can arise through modification of cell-wall architecture rather than changes in arabinoxylan biosynthetic flux. The importance of cell-wall peroxidases in determining AX extractability is further supported by the recent identification of PER2A, the chromosome 6A homoeologue of the tandem copy of PER1, which was independently associated with WE-AX by Li et al. (2026). PER2A exhibits a developmental expression profile similar to that of PER1 in the starchy endosperm (Mitchell et al., 2026), suggesting that these homoeologous peroxidases perform conserved biological functions. We did not detect a significant association at the PER2A locus, most likely because the causal variant reported by Li et al. (2026) is absent from the Elite Fibre Panel analysed here. Interestingly, although additional class III peroxidases were identified within the chromosome 7A QTL interval, these genes, unlike PER1 and PER2A, are not commelinid-specific. Because feruloylated arabinoxylans are a defining feature of commelinid primary cell walls, this evolutionary distinction suggests that the chromosome 7A peroxidases are less likely to function as specialised regulators of ferulate-mediated arabinoxylan cross-linking.

Unlike the relatively simple PER1 locus, the major chromosome 1B QTL encompasses several biologically plausible candidate genes and probably represents a more complex regulatory region. Previous studies have consistently shown that this locus influences both total arabinoxylan and water-extractable arabinoxylan (Lovegrove et al., 2020; Ibba et al., 2021), suggesting that it affects polysaccharide accumulation rather than solubility alone. Among the prioritised candidates, the grass-specific xyloglucan endotransglucosylase/hydrolase (XTH/GH16) represents an attractive candidate because XTH proteins regulate cell-wall remodelling through cleavage and rejoining of xyloglucan chains, thereby influencing wall architecture and polysaccharide organisation. Although *TraesCS1B02G424500* showed its highest transcript abundance during the early stages of endosperm development, XTH-mediated cell-wall remodelling during this period could influence the assembly of the nascent primary cell wall and the subsequent deposition and organisation of arabinoxylan. XTH enzymes catalyse the cleavage and religation of xyloglucan chains and are recognised as key regulators of primary cell-wall restructuring during plant growth and development (Rose et al., 2002). Furthermore, while most characterised XTHs use xyloglucan as both donor and acceptor substrates, some members of the family exhibit broader transglycosylase activity towards non-canonical hemicellulose substrates, suggesting that XTH-mediated remodelling may influence interactions among cell-wall polysaccharides (Maris et al., 2011). Although XTH proteins have not been directly implicated in wheat arabinoxylan biosynthesis or modification, their established role in primary cell-wall remodelling, together with the early developmental expression of *TraesCS1B02G424500*, makes this gene a strong biological candidate underlying the chromosome 1B QTL. Additional candidates within this interval include a fasciclin-like arabinogalactan protein involved in cell-wall organisation and an endosperm-specific class III peroxidase that could contribute to oxidative modification of cell-wall phenolics. Together, these observations suggest that the chromosome 1B locus may regulate multiple aspects of cell-wall assembly rather than a single biochemical pathway.

Beyond the two major loci, the additional QTLs identified in this study reveal several complementary biological processes that may contribute to natural variation in WE-AX. The chromosome 5B locus containing UTP-glucose-1-phosphate uridylyltransferase (UGPase) represents a particularly compelling candidate because UGPase produces UDP-glucose, the central precursor for nucleotide sugars that ultimately generate UDP-xylose and UDP-arabinose, the activated sugar donors required for arabinoxylan biosynthesis (Kleczkowski et al., 2004). Rather than influencing arabinoxylan structure directly, this candidate is likely to regulate substrate availability for cell-wall polysaccharide synthesis. In contrast, the second chromosome 5B locus may influence WE-AX through regulatory rather than biosynthetic mechanisms. The highest-priority candidate, a commelinid-specific pentatricopeptide repeat (PPR)-containing protein, belongs to a family traditionally associated with organellar RNA processing, but increasing evidence demonstrates that specific PPR proteins participate in cell-wall integrity signalling and maintenance through mitochondria-to-nucleus communication (Hu et al., 2016; Hofmann, 2016). Consequently, this locus may influence arabinoxylan accumulation indirectly through regulatory networks controlling cell-wall development. Although additional candidates at this locus, including a SAM-dependent methyltransferase and an FAD/NAD(P)-binding oxidoreductase, have no direct experimental connection to arabinoxylan metabolism, their known roles in methyl-group metabolism, redox homeostasis and phenylpropanoid metabolism make them biologically plausible candidates deserving further investigation.

The chromosome 7A interval identified several candidates with potential roles in hemicellulose modification and cell-wall maturation. Particularly noteworthy are the TBL/DUF231 proteins, members of a family experimentally demonstrated to function as xylan O-acetyltransferases in both Arabidopsis and rice (Gao et al., 2017). Altered xylan acetylation influences xylan interactions with cellulose and other wall polymers and therefore represents a plausible mechanism affecting arabinoxylan extractability. The co-occurrence within the same interval of a nucleotide-sugar transporter, a xyloglucan endotransglucosylase, a cationic peroxidase and glycosyltransferase-related genes further strengthens the biological plausibility of this region as a regulator of grass cell-wall composition. In contrast, the second chromosome 7A association (AX-643821508) could not be resolved to candidate genes because no neighbouring SNPs were in significant linkage disequilibrium with the lead marker, resulting in an LD interval that collapsed to the position of the associated SNP. This may reflect limited local marker density, structural variation absent from the reference genome, or a causal variant located within an unannotated regulatory region.

The chromosome 3A locus represents a contrasting example of a genetically simple interval, containing only a single annotated gene encoding a RING/U-box protein. Members of this family function primarily as E3 ubiquitin ligases that regulate protein turnover during plant development and stress responses. Although no direct evidence currently links this gene family to arabinoxylan biosynthesis, the presence of only one annotated gene within the LD interval makes TraesCS3A02G439700 an attractive target for future functional validation.

Developmental expression profiles provided an additional independent line of evidence supporting candidate-gene prioritisation (Fig. 7). Previous transcriptomic analyses of developing wheat endosperm have shown that the period between approximately 10 and 25 days after anthesis corresponds to active cell-wall deposition, when expression of genes involved in arabinoxylan biosynthesis reaches its maximum (Pellny et al., 2012). Consistent with this developmental programme, UGPase exhibited exceptionally high expression throughout this period, supporting its proposed role in supplying nucleotide-sugar precursors for polysaccharide synthesis. Likewise, PER1 showed sustained expression during mid-to-late grain development, consistent with post-depositional oxidative cross-linking accompanying cell-wall maturation. In contrast, the chromosome 3A RING/U-box protein displayed its highest expression at 4 dpa before declining steadily, suggesting a role during early endosperm differentiation rather than during active arabinoxylan deposition. The chromosome 7A TBL candidate exhibited a distinct expression peak around 21-25 dpa, coinciding with the transition towards grain maturation, consistent with the proposed role of TBL proteins in post-synthetic modification of xylan. These contrasting expression profiles support the view that the identified loci influence different stages of cell-wall development rather than acting within a single biosynthetic pathway.

The evolutionary conservation analysis provided an additional and independent layer of evidence for candidate prioritisation. Several of the highest-ranked candidates, including PER1, the chromosome 7A TBL protein and the chromosome 5B PPR protein, were classified as commelinid- or grass-specific using the Universal Grass Gene Database (Mitchell, 2025). This observation is biologically meaningful because feruloylated arabinoxylan is a defining feature of commelinid monocot primary cell walls and is absent from dicot species (Schendel et al., 2016; Terrett and Dupree, 2019). Consequently, evolutionary specificity provides complementary support that these genes have evolved specialised functions associated with grass cell-wall biology, strengthening their candidacy beyond positional and expression evidence alone.

Taken together, the candidate genes identified in this study represent multiple distinct biological processes that collectively regulate soluble arabinoxylan accumulation. Rather than identifying genes acting within a single pathway, our results suggest that natural variation in WE-AX reflects coordinated genetic variation affecting precursor supply (UGPase), cell-wall assembly/remodelling (XTH), polysaccharide modification (TBL proteins), oxidative cross-linking (PER1), and regulatory processes governing cell-wall development (PPR and RING/U-box proteins). This mechanistic diversity provides a coherent explanation for the largely additive genetic effects observed across loci and highlights multiple entry points through which breeding can manipulate arabinoxylan content.

The additive effects of favourable alleles have important practical implications for wheat improvement. Genotypes carrying progressively larger numbers of high-fibre alleles showed stepwise increases in WE-AX content, demonstrating that these loci contribute largely independently to phenotypic variation. Because favourable alleles remain segregating within elite germplasm, substantial improvements in soluble fibre content should be achievable through marker-assisted selection or genomic selection without introducing exotic germplasm and the associated risk of linkage drag. The development of diagnostic KASP markers for the newly identified loci should further accelerate deployment of these alleles in commercial breeding programmes.

Although the present study provides strong evidence for several high-confidence candidate genes, further work will be required to establish gene-level causality for most loci. Fine mapping, allele-specific expression analyses, variant-effect prediction, and functional validation using genome editing or mutant resources will be essential to distinguish causal genes from linked candidates within the larger LD intervals. Integrating GWAS with transcriptomics, comparative genomics and biochemical analyses, as demonstrated here, provides a powerful framework for accelerating candidate-gene discovery and prioritisation in complex polygenic traits. The genetic architecture defined in this study provides a practical framework for breeding wheat with enhanced soluble dietary fibre. Because water-extractable arabinoxylan in wholemeal flour is highly correlated with that in white flour (Lovegrove et al., 2020), the loci identified here are directly relevant to improving the nutritional quality of the wheat products most widely consumed worldwide.

Given the well-established associations between dietary fibre and improved gut health, glycaemic regulation, and reduced risk of chronic metabolic disease, even modest genetic increases in WE-AX content in widely consumed white flour products could translate into substantial public-health benefits at the population scale. By directly linking genetic variation to a nutritionally relevant trait in breeding-relevant material, this study bridges the gap between genomics and health-oriented crop improvement and highlights the potential of wheat breeding as a scalable and sustainable intervention for improving dietary fibre intake.

## Acknowledgements

The authors sincerely thank the members of the Elite Fibre Panel (EFP) Breeders’ Consortium for generously providing the elite wheat germplasm used in this study: Simon Berry (Limagrain UK Ltd., Suffolk, UK), Alastair Nash (RAGT Seeds Ltd., Essex, UK), Andrew Creasy (Saaten Union UK Ltd., Cambridgeshire, UK), and Bill Angus (F1 Seeds Ltd., Suffolk, UK). Their continued support of collaborative research and willingness to share elite breeding material were instrumental to the success of this study.

## Author contributions

AKA curated the phenotypic and genotypic datasets, performed the genome-wide association analyses and other genetic and statistical analyses, identified and prioritised candidate genes, generated all figures, and wrote the first draft of the manuscript. AL, OK, and AP performed the wholemeal arabinoxylan analyses. ML-W coordinated germplasm collection from wheat breeders and prepared DNA samples for genotyping. RACM contributed to candidate gene prioritisation and interpretation. JB developed and provided the chromosome 1B KASP markers. SG, PRS, and AL conceived and designed the study and secured funding. All authors contributed to data interpretation, critically revised the manuscript, and approved the final version of the manuscript.

## Data availability

All data supporting the findings of this study are provided within the article and its supplementary materials. The datasets generated and analysed during the current study are available from the corresponding author on reasonable request.

## Supplementary Information

Additional Supporting Information may be found online in the Supporting Information section at the end of the article.

**Fig. S1** Quality control summaries of the filtered SNP dataset used for genome-wide association analysis of the Elite Fibre Panel.

**Fig. S2** Consistency of principal component correction across GWAS models.

**Fig. S3** Heatmap of the pairwise kinship matrix of the Elite Fibre Panel.

**Fig. S4** High-fibre allele frequencies across the Elite Fibre Panel (EFP) and other germplasms.

**Fig. S5** Genome-wide linkage disequilibrium (LD) patterns of the Elite Fibre Panel.

**Fig. S6** Chromosome-wide linkage disequilibrium (LD) heatmaps for the five wheat chromosomes carrying significant WE-AX quantitative trait loci (QTL).

**Fig. S7** Chromosome-wide linkage disequilibrium (LD) decay for chromosomes harbouring significant WE-AX loci.

**Table S1** Detailed information of the Elite Fibre Panel (EFP) lines

**Table S2** Wholemeal flour arabinoxylan (xylose) content of the Elite Fibre Panel (EFP) across the Limagrain and RAGT field trial sites and their across-environment mean (Average).

**Table S3** Detailed information on the KASP markers used to extend the Elite Fibre Panel (EFP) genotypic data used in the GWAS analysis.

**Table S4** Details of marker filtering for genotyping the Elite Fibre Panel (EFP).

**Table S5** Single-bit HapMap-formatted genotypic matrix of the Elite Fibre Panel (EFP), generated from Axiom TaNG1.1 SNP array and KASP genotyping after quality control filtering, and used for GAPIT-based genome-wide association analyses.

**Table S6** Genome-wide significant marker-trait associations for wholemeal flour arabinoxylan (xylose) content identified using BLINK, FarmCPU, and MLMM models across the Limagrain and RAGT field trial sites and their across-environment mean.

**Table S7** Genotypes at the seven high-fibre-associated SNP markers across three wheat genetic resources.

**Table S8** Metadata of the publicly available wheat starchy endosperm RNA-seq datasets used to evaluate the developmental expression profiles of candidate genes identified by GWAS.

**Table S9** List of genes identified within LD-defined QTL intervals associated with wholemeal flour water-extractable arabinoxylan (WE-AX) content in the Elite Fibre Panel (EFP).

