## Supplementary Figures for "Genetic architecture of soluble arabinoxylan fibre in elite genotypes of bread wheat revealed by genome-wide association analysis"

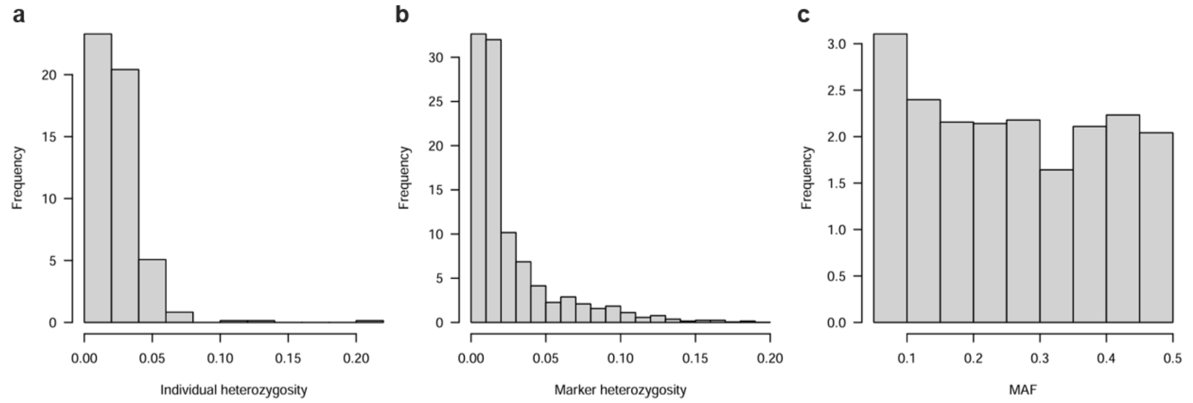


**Fig. S1 Quality control summaries of the filtered SNP dataset used for genome-wide association analysis of the EFP.** (**a**) Distribution of individual heterozygosity among the 365 wheat accessions. (**b**) Distribution of marker heterozygosity across the 6,791 SNP markers retained after quality filtering. (**c**) Distribution of minor allele frequencies (MAF) for the filtered SNP markers. SNPs with MAF < 0.05 and markers with high levels of missing data or heterozygosity were excluded before GWAS to reduce spurious associations and improve the robustness of downstream analyses. The approximately uniform distribution of common alleles demonstrates that the filtering strategy retained sufficient allele frequency diversity for GWAS while excluding rare variants with limited statistical power.

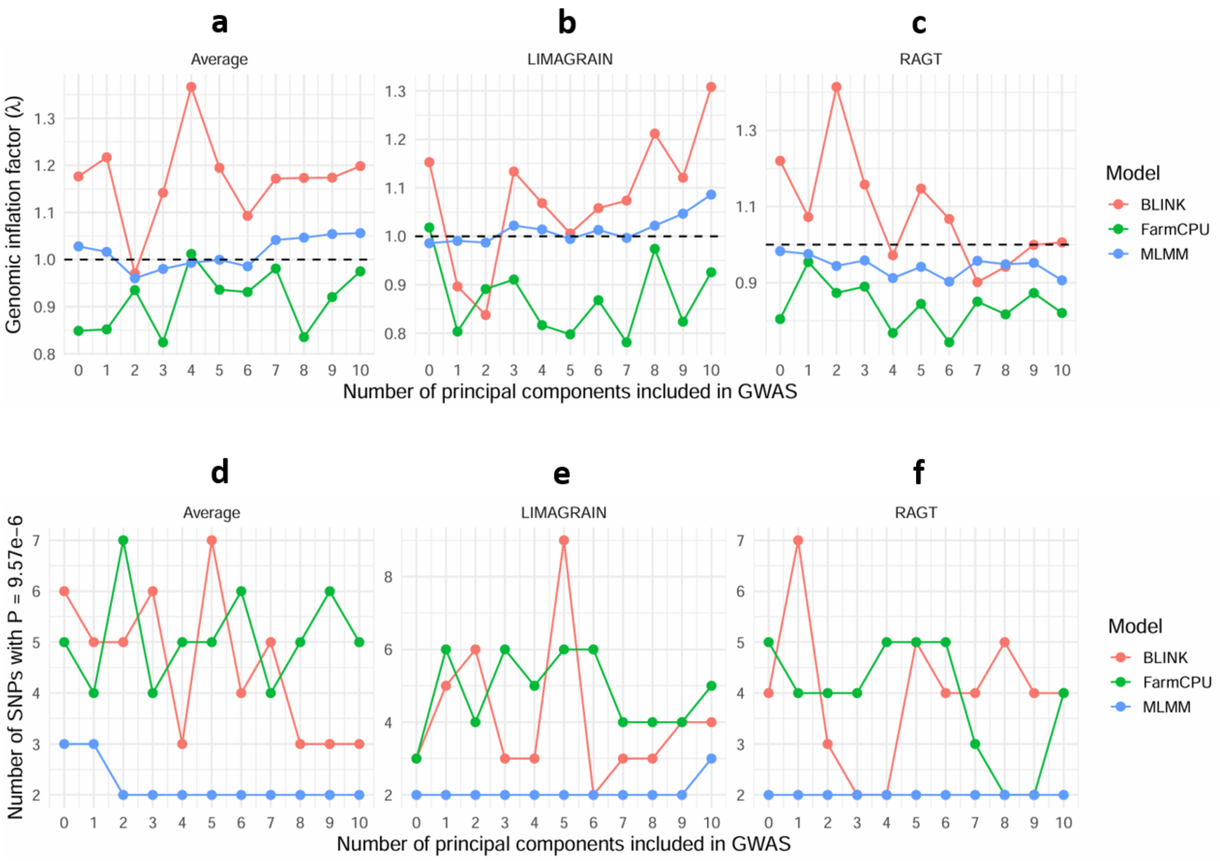


**Fig. S2 Consistency of principal component correction across GWAS models.** Genomic inflation factor (λ) and the number of significant marker–trait associations obtained using the three best-performing GWAS models (BLINK, FarmCPU, and MLMM) following inclusion of 0-10 PCs as fixed covariates. Panels (**a - c**) show the genomic inflation factor (λ) for the Average, Limagrain, and RAGT phenotypic datasets, respectively. The dashed horizontal line indicates the expected value under the null hypothesis (λ = 1). Across all three datasets, genomic inflation remained close to 1 regardless of the number of PCs included, although BLINK generally produced slightly higher λ values, whereas FarmCPU tended to produce slightly lower values and MLMM remained consistently close to unity. No systematic improvement in model calibration was observed with increasing numbers of principal components. Panels (**d - f**) show the number of genome-wide significant associations ($P \leq 9.57 \times 10^{-6}$) detected by BLINK, FarmCPU, and MLMM for the Average, Limagrain, and RAGT datasets, respectively. The number of significant loci was largely stable across the tested PC values, particularly for MLMM, while BLINK and FarmCPU showed only modest fluctuations without any consistent trend. Collectively, these analyses demonstrate that the inclusion of principal components had minimal influence on genomic inflation or the detection of significant marker–trait associations across the three GWAS models, providing independent support for the use of **PCA.total = 0** in the final GWAS analyses.

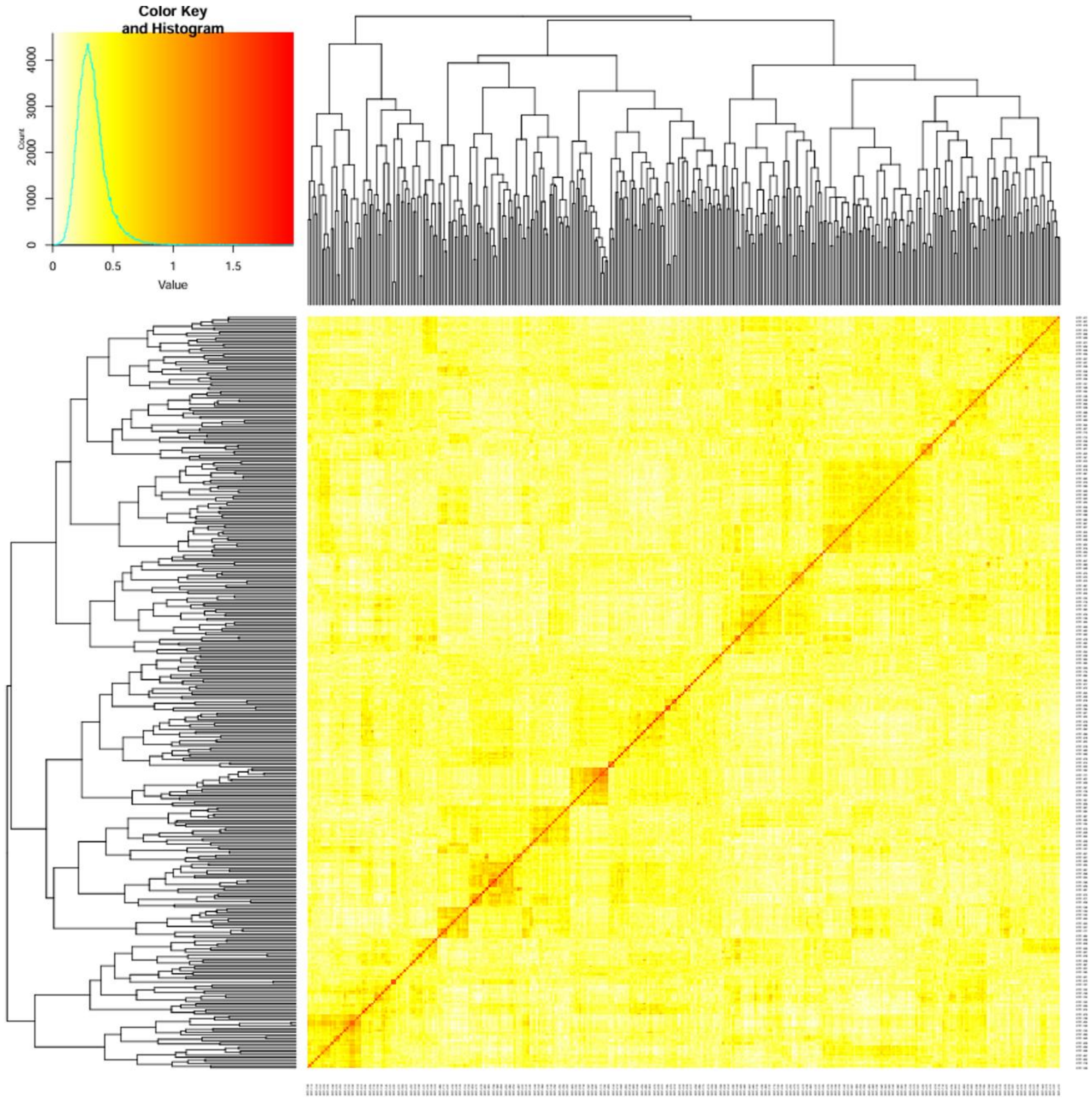


**Fig. S3 Heatmap of the pairwise kinship matrix of the Elite Fibre Panel.** Pairwise genomic kinship matrix was calculated using the Zhang method in GAPIT from 6,791 high-quality SNP markers (MAF ≥ 0.05). The heatmap illustrates the realised genetic relationships among the 365 accessions, with warmer colours (red) indicating higher kinship coefficients. Hierarchical clustering identifies groups of genetically similar accessions and reveals patterns of pairwise relatedness within the panel. The predominance of low off-diagonal kinship coefficients indicates limited relatedness among accessions, although several small clusters of closely related genotypes were detected. Incorporating the kinship matrix into the mixed-model analyses, therefore, helped control for cryptic relatedness and reduce false-positive associations.

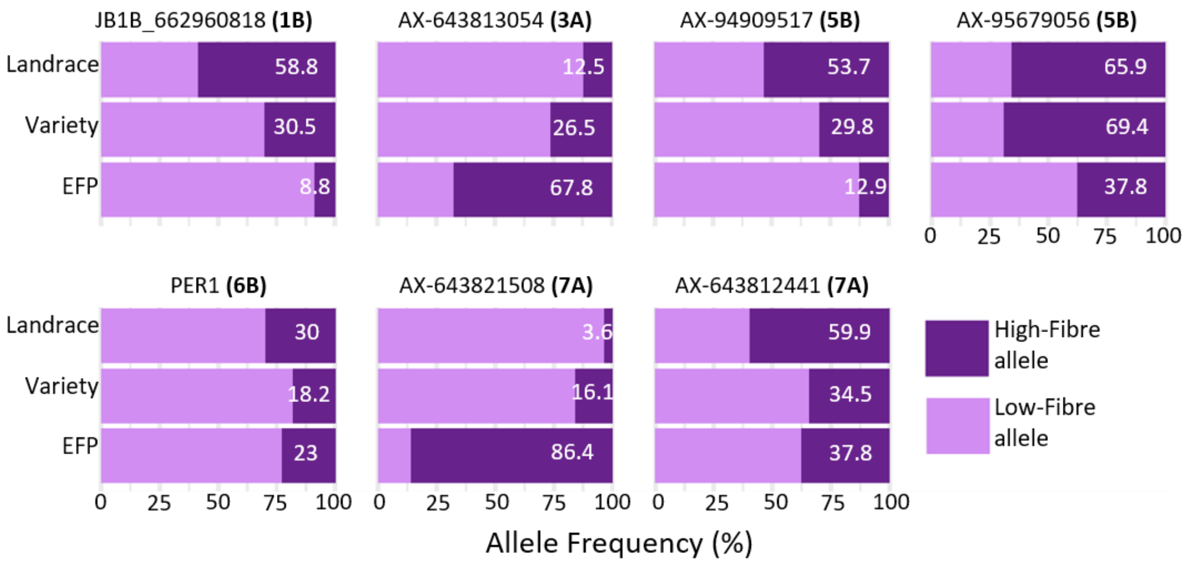


**Fig. S4 High-fibre allele frequencies across the Elite Fibre Panel (EFP) and other germplasms.** Stacked bar plots showing the frequency (%) of the high-fibre allele (dark purple) and the low-fibre allele (light purple) at seven significant GWAS-associated SNP markers (JB1B_662960818, AX-643813054, AX-94909517, AX-95679056, PER1, AX-643821508, and AX-643812441) across three wheat genetic resources: the Elite Fibre Panel (EFP), modern varieties (Variety), and Watkins landraces (Landrace). Allele frequencies were calculated within each genetic resource based on genotype calls generated in this study from Axiom array genotyping of the EFP and from SNP calls derived from whole-genome resequencing of 827 Watkins landraces and 224 modern varieties (Cheng et al., 2024). Numbers on the bars indicate the frequency (%) of the high-fibre allele.

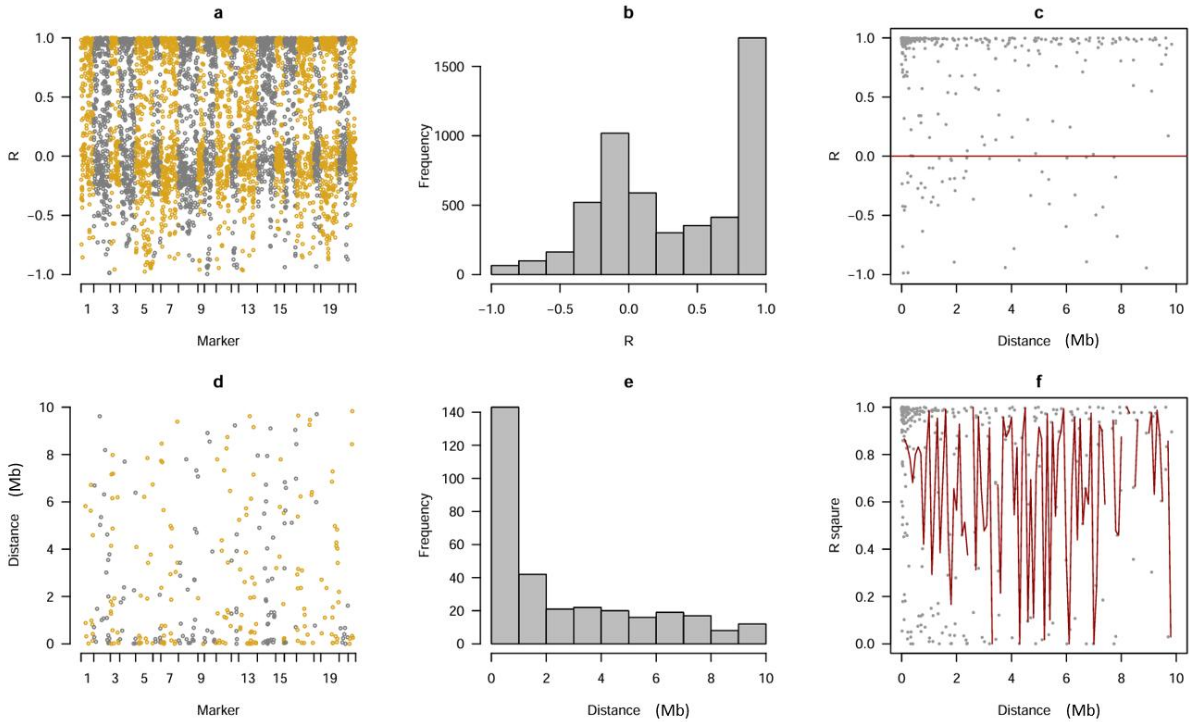


**Fig. S5 Genome-wide linkage disequilibrium (LD) patterns of the Elite Fibre Panel (EFP).** LD analysis performed by GAPIT using the 6,791 high-quality SNP markers and used for GWAS of the EFP. (**a**) Pairwise correlation coefficients (r) between neighbouring SNP markers across the 21 wheat chromosomes. (**b**) Frequency distribution of pairwise marker correlations. (**c**) Relationship between marker correlation (r) and physical distance between adjacent markers. (**d**) Physical distances separating neighbouring SNP markers across the genome. (**e**) Distribution of inter-marker distances. (**f**) Relationship between LD ($r^2$) and physical distance, with the red curve representing the locally weighted regression (LOESS) fit. Overall, the marker set provided broad genome-wide coverage, with most neighbouring markers separated by less than 2 Mb. Although marker density varied across the genome, the coverage was sufficient to capture common haplotypes within the EFP and support genome-wide association mapping. As expected, linkage disequilibrium generally declined with increasing physical distance, although substantial variation in $r^2$ was observed across the genome, reflecting local differences in recombination history, marker density, and haplotype structure. The persistence of moderate to high LD over short genomic intervals indicates that the marker density was sufficient to capture most common haplotypes within the Elite Fibre Panel and therefore provided adequate resolution for genome-wide association mapping.

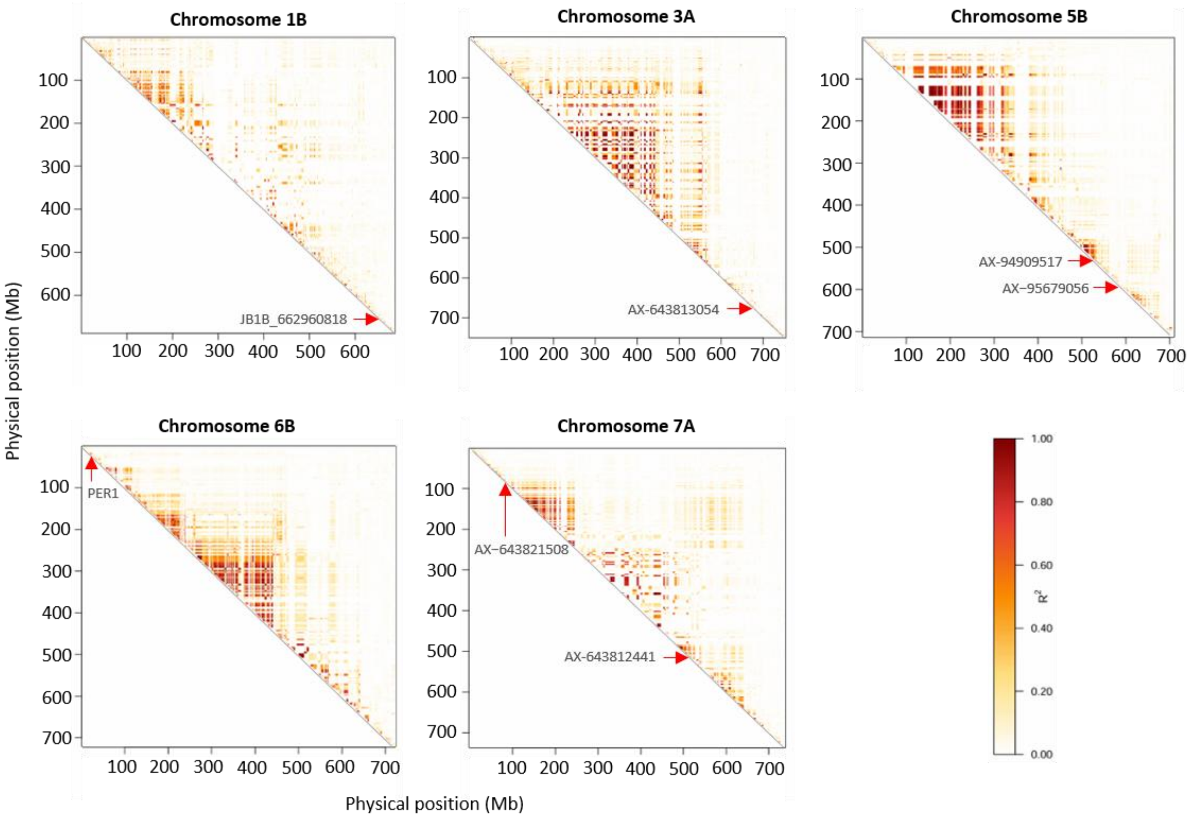


**Fig. S6 Chromosome-wide linkage disequilibrium (LD) heatmaps for the five wheat chromosomes carrying significant WE-AX quantitative trait loci (QTL).** Pairwise linkage disequilibrium (LD) was calculated among all SNPs located on chromosomes 1B, 3A, 5B, 6B and 7A using the squared correlation coefficient ($R^2$). SNPs were ordered according to their physical positions on the reference genome, and only the upper triangular portion of each symmetric LD matrix is displayed. Colour intensity represents the magnitude of LD, ranging from white ($R^2 \approx 0$) to dark red ($R^2 = 1$). Red arrows indicate the physical positions of the lead SNPs identified by GWAS (JB1B_662960818 on chromosome 1B, AX-643813054 on chromosome 3A, AX-94909517 and AX-95679056 on chromosome 5B, PER1 on chromosome 6B, and AX-643821508 and AX-643812441 on chromosome 7A). Localised blocks of high LD indicate genomic regions where neighbouring markers exhibit strong allelic association owing to limited historical recombination. In contrast, the predominance of low $R^2$ values across each chromosome reflects extensive historical recombination and generally rapid LD decay within the EFP. The heterogeneous distribution of LD across chromosomes illustrates variation in local recombination history and haplotype structure surrounding the detected QTL.

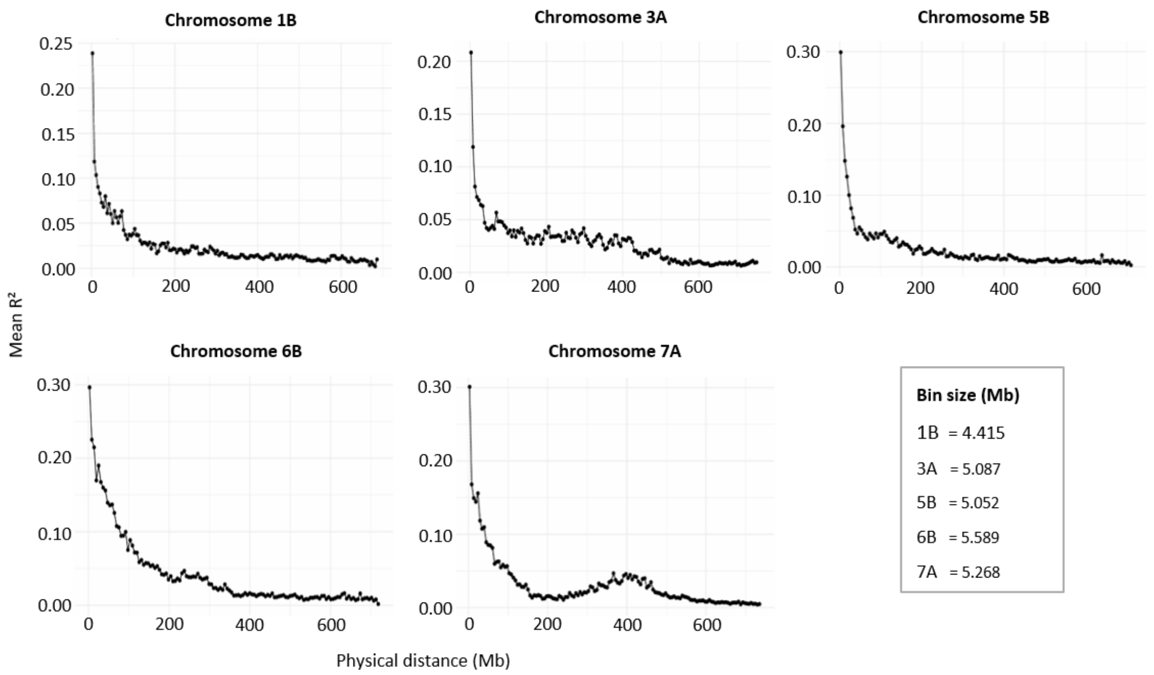


**Fig. S7 Chromosome-wide linkage disequilibrium (LD) decay for chromosomes harbouring significant WE-AX loci.** Pairwise linkage disequilibrium (LD) was calculated among all SNP pairs located on chromosomes 1B, 3A, 5B, 6B, and 7A, which harbour significant WE-AX quantitative trait loci (QTL). LD was estimated as the squared correlation coefficient ($R^2$) between SNP pairs and grouped into adaptive physical-distance bins containing a minimum of 30 SNP pairs. Mean LD ($R^2$) was calculated for each bin and plotted against the midpoint of the corresponding physical-distance interval. All chromosomes exhibited the expected decline in LD with increasing physical distance, reflecting the progressive breakdown of marker associations through historical recombination. Although all chromosomes showed rapid initial LD decay, the rate and shape of the decay curves differed among chromosomes, indicating variation in local recombination history, haplotype structure, and marker density. These chromosome-wide LD patterns provide a general estimate of the extent of linkage disequilibrium within the Elite Fibre Panel and complement the marker-specific LD analyses used to define QTL intervals (Fig. 6).
